# Epigenetic mechanisms mediating cell state transitions in chondrocytes

**DOI:** 10.1101/2020.10.26.319822

**Authors:** Manuela Wuelling, Christoph Neu, Andrea M. Thiesen, Simo Kitanovski, Yingying Cao, Anja Lange, Astrid M. Westendorf, Daniel Hoffmann, Andrea Vortkamp

**Author notes:** These authors contributed equally to this work (Manuela Wuelling, Christoph Neu).

## Abstract

Epigenetic modifications play critical roles in regulating cell lineage differentiation, but the epigenetic mechanisms guiding specific differentiation steps within a cell lineage have rarely been investigated. To decipher such mechanisms, we used the defined transition from proliferating (PC) into hypertrophic chondrocytes (HC) during endochondral ossification as a model. We established a map of activating and repressive histone modifications for each cell type. ChromHMM state transition analysis and Pareto-based integration of differential levels of mRNA and epigenetic marks revealed that differentiation associated gene repression is initiated by the addition of H3K27me3 to promoters still carrying substantial levels of activating marks. Moreover, the integrative analysis identified genes specifically expressed in cells undergoing the transition into hypertrophy.

Investigation of enhancer profiles detected surprising differences in enhancer number, location, and transcription factor binding sites between the two closely related cell types. Furthermore, cell type-specific upregulation of gene expression was associated with a shift from low to high H3K27ac decoration. Pathway analysis identified PC-specific enhancers associated with chondrogenic genes, while HC-specific enhancers mainly control metabolic pathways linking epigenetic signature to biological functions.

## Introduction

In the last decade, the role of epigenetic modifications in regulating tissue differentiation and development has been intensively studied. Most of these studies have, however, focused on the differentiation of distinct cell lineages from embryonic or the mesenchymal stem cells or the comparison of distinct tissues (1, 2) but the epigenetic modifications regulating changes in gene expression at discrete differentiation steps within a specific cell lineage are less well understood. To receive insight into such epigenetic mechanisms we have investigated the transition of proliferating into hypertrophic chondrocytes during endochondral ossification as a representative differentiation step. The switch between the two differentiation states takes place at a precise morphological position and is tightly controlled by numerous transcription factors, signaling pathways and epigenetic modifiers (3–6) allowing to relate differences in the epigenetic profile to gene function.

Endochondral ossification is initiated by the formation of a Sox9 and Collagen type 2a1 (Col2) expressing condensation of proliferating chondrocytes (PC). Once these templates of the later bones have reached a critical size, PC in their center exit the cell cycle and differentiate into hypertrophic chondrocytes (HC), which express Collagen type 10a1 (Col10) instead of Col2 and mineralize their extracellular matrix (ECM). This transition in critical in balancing growth and stability of the later skeletal elements as HC are subsequently replaced by ossified bone tissue.

Epigenetic modifications relevant for the formation of distinct tissues include the acetylation and methylation of specific histone residues. While histone acetylation results in an opening of the chromatin structure and, consequently, the activation of gene expression, their deacetylation leads to a dense chromatin structure and gene silencing. Acetylated histones are localized at distinct regulatory regions, as for example acetylated lysine 9 of histone 3 (H3K9ac), which demarcates active promoters, while acetylated lysine 27 (H3K27ac) marks active promoters and enhancers. In contrast to acetylation, histone methylation elicits positive or negative effects on gene expression depending on the position of the methylated residue. Trimethylation of lysine 4 of histone 3 (H3K4me3) defines the promoter of active genes and genes primed for expression (7). Trimethylation of lysine 36 of histone H3 (H3K36me3) is associated with actively transcribed genes (8). In contrast, trimethylation of histone H3 at lysine 9 (H3K9me3) or lysine 27 (H3K27me3) (9, 10) leads to gene repression. Besides different combinations of activating marks at promoter regions, the combination of activating H3K4me3 and repressive H3K27me3, the so-called bivalent mark, is found at promoters of many lineage-specific genes in embryonic stem cells and has been associated with pluripotency (11). During differentiation, genes carrying this bivalent mark can either gain additional activating histone marks and lose H3K27me3 to initiate transcription or lose H3K4me3 leading to gene repression (12, 13). Recently, another bivalent chromatin state, a combination of H3K36me3 and H3K9me3, has been described on weakly transcribed genes (14, 15).

In chondrocyte, first studies investigated epigenetic marks associated with binding of the transcription factor Sox9 in postnatal rib chondrocytes (16) and rat chondrosarcoma cells (17). More recently, the analysis of open chromatin in mixed growth plate chondrocytes by ATAC-seq identified specific enhancers modulating skeletal heights (18) and osteoarthrosis susceptibility, but a systematic investigation of the epigenetic mechanisms controlling distinct steps of chondrocyte differentiation *in vivo* is still lacking.

In this study, we have used the transition from PC to HC to receive insight into the epigenetic alterations linked to a switch in cell state and have correlated them to differences in gene expression. By establishing an unbiased epigenetic profile of activating and repressive marks we identified the addition of H3K27me3 to promoter regions still carrying activating marks as the gene repression inducing event. The analysis of H3K27ac occupancy revealed an unexpected difference in enhancer usage and a rapid accumulation and decline with changes in gene expression. Besides this mechanistic insight into gene regulation our study identified metabolism as the main biological function of hypertrophic chondrocytes.

## Results

### Cell type-specific epigenetic profiles of chondrocytes

To identify epigenetic changes linked to the differentiation of PC into HC, we isolated the respective cells from transgenic mice expressing YFP under the control of the Collagen Type 2a1 (Col2-Cre;R26R-YFP) or the Collagen Type 10 (Col10-Cre;R26R-YFP) promoter by fluorescence-activated cell sorting (FACS). Flow cytometry of cells isolated from E13.5 Col2-Cre;R26R-YFP skeletons, a time point before hypertrophic differentiation takes place, detected 15% to 20% YFP-positive cells, which were enriched to about 90% by FACS. At E15.5, only 1% of the skeletal cells of Col10-Cre;R26R-YFP mice were YFP positive. These were enriched to approximately 80% YFP-positive cells (Fig. 1A-B).

**Figure 1:**
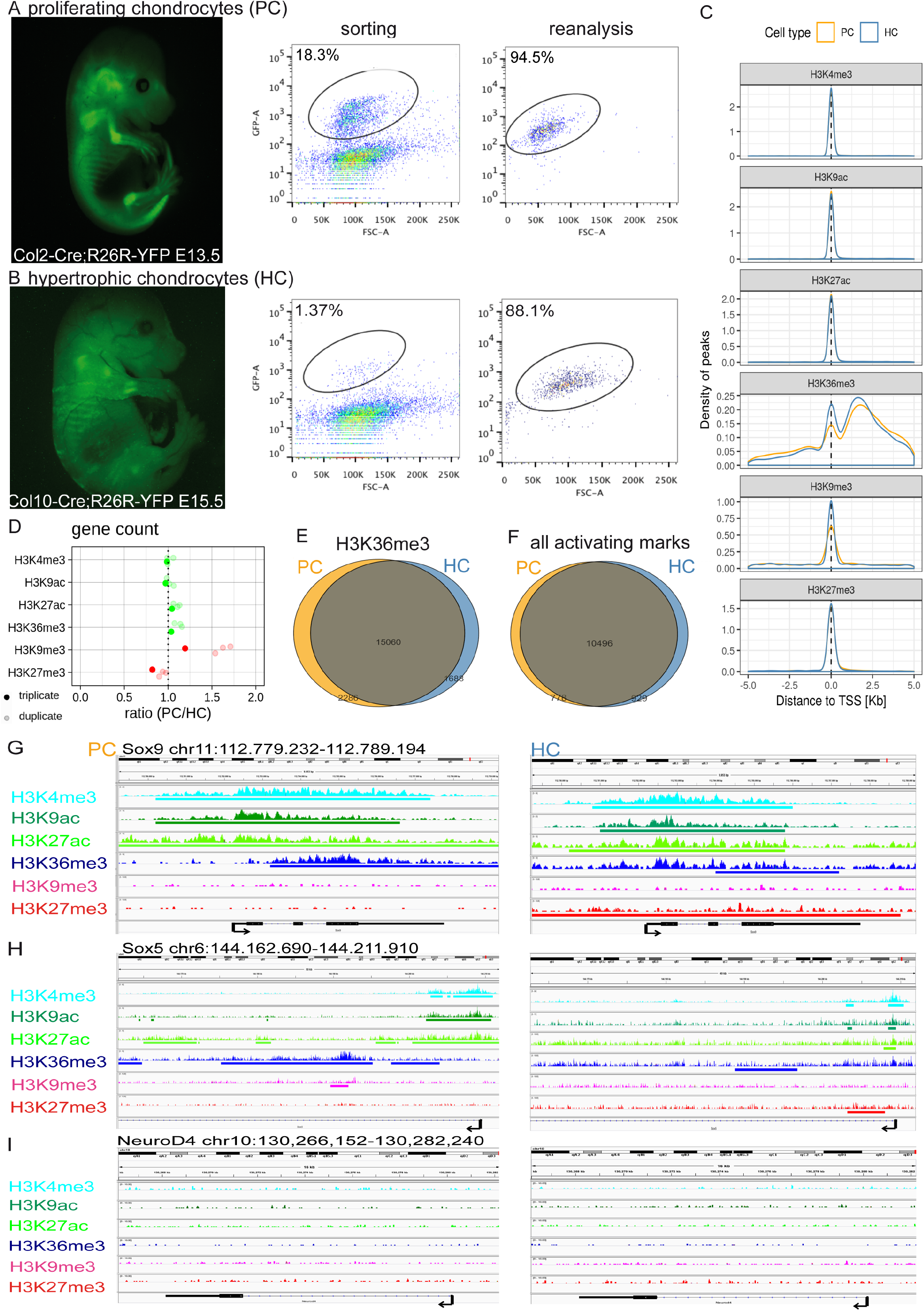
Epigenetic profiling of PC and HC. (A-B) Fluorescent images and flow cytometry of cells isolated from E13.5 Col2-Cre;R26R-YFP (A) and E15.5 Col10-Cre;R26R-YFP (B) mice prior to (sorting) and after cell sorting (reanalysis), YFP positive populations (black ellipse). (C) Peak density of the histone marks around the TSS (+/−5 kb, left) in PC (orange) and HC (blue). The activating histone marks H3K4me3, H3K9ac and H3K27ac cluster at the TSS. (D) Ratio (PC/HC) of the genes marked by the indicated histone mark, activating marks (green) repressive marks (red). (E-F) Venn diagram of genes carrying H3K36me3 (E) and the activating histone marks H3K4me3, H3K9ac, H3K27ac and H3K36me3 (F). (G-I) Read and peak coverages of histone marks in the genomic regions of Sox9 (G), Sox5 (H) and NeuroD4 (I) visualized in IGV; TSS (black arrow).

To detect differences in the epigenetic profile of the two chondrocyte populations, we investigated the distribution of the activating promoter marks H3K4me3 and H3K9ac, the promoter and enhancer mark H3K27ac, the transcription-associated mark H3K36me3 and the repressive marks H3K27me3 and H3K9me3 by ChIP-seq (Suppl.Fig. 1A). Due to the restricted number of HC per embryo, the ChIP-seq experiments were performed in biological triplicates for PC and in duplicates for HC. Principal component analysis and hierarchical clustering revealed clustering according to distinct histone marks (Suppl.Fig. 1B-C) and a high similarity between the activating and repressive modifications. H3K36me3 showed a similar correlation with both groups (Suppl.Fig. 1C), likely reflecting the potential overlap with activating and repressive marks (14). Genomic regions enriched for each chromatin mark (Suppl.Table 1) were identified by hiddenDomains (19). To exclude that different replicate numbers in PC and HC affect the analysis, we compared peak coverage and associated gene number of the PC datasets pairwisely but did not detect major differences between the PC triplicate and the different subsets of PC duplicates (Fig. 1D, Suppl.Fig 1D-E). We compiled a dataset (conserved dataset) of all peaks that were conserved throughout all replicates and used this dataset for further analyses. Peak distribution analysis detected the expected enrichment of activating promoter marks and the repressive modification H3K27me3 at the transcription start site (TSS +/−1 kb), whereas H3K36me3 and H3K9me3 were distributed throughout the genome (Fig. 1C). Comparison of peak coverage and gene number revealed a similar distribution of most modifications in PC and HC and a slightly increased number of H3K27me3 peaks in HC (Fig. 1D, Suppl.Fig. 1D-G).

Analysis of genes marked as transcribed by H3K36me3 identified 2286 genes specifically expressed in PC and 1688 in HC, while 15060 genes were expressed in both cell types, reflecting the common chondrogenic lineage (Fig. 1E). A large fraction of these genes was additionally marked by the three activating promoter marks (Fig. 1F). Evaluation of genes known to be downregulated in HC, like Sox9 and Sox5 (17, 20) revealed a high coverage of activating promoter marks at the TSS and H3K36me3 on the gene body in PC, while repressive marks were absent. Interestingly, in HC, these genes gain the repressive mark H3K27me3 especially at the promoter region but remain decorated with reduced levels of activating marks (Fig. 1G-H), indicating that gene repression is initiated by the addition of H3K27me3 in regions still marked as active. In contrast, genes not expressed in chondrocytes, like NeuroD4, did not carry repressive marks in either cell type, making an unspecific increase in histone methylation during differentiation unlikely (Fig. 1I).

### Genes marked as active gain H3K27me3 during differentiation

In order to identify epigenetic changes linked to differentiation in an unbiased approach, we defined 15 chromatin states carrying distinct combinations of epigenetic marks using ChromHMM (Fig. 2A, Suppl.Fig.2A-B, Suppl.Fig. 3 A-C). Based on the combination of histone marks the chromatin states were classified as activating (states 8, 9, 11, 13), enhancer (states 5, 7, 10), expressed (states 3 (14, 15), 4), repressive (states 1, 2, 14) and empty (state 6). In addition, ChromHMM identified two chromatin states with a combination of H3K27me3 and activating promoter marks (state 13) or H3K36me3 (state 15), here referred to as repressed-active states (Fig. 2A). In the same cell type, transitions between states along the genome occurred with highest probability between functionally related states or into the empty state 6 (Suppl.Fig. 2C). Functional assignment revealed the expected, region-specific distribution of states with promoter associated states 8 to 13 enriched at the TSS and CpG islands. The enhancer states 5, 7 and 10 colocalized with published enhancers extracted from the Vista and Fantom databases (Suppl.Fig. 2D). To confirm that the identified ChromHMM states represent histone combinations frequently occurring during development, we included epigenetic data from E11.5-E15.5 limb bud and brain tissue into the model. Besides the detection of more promoter states, we did not find major differences in the generalized model. The most interesting difference was the loss of the repressed-active state 15, indicating that the combination of activating and repressive histone marks was overrepresented in our dataset (Suppl.Fig. 3D).

**Figure 2:**
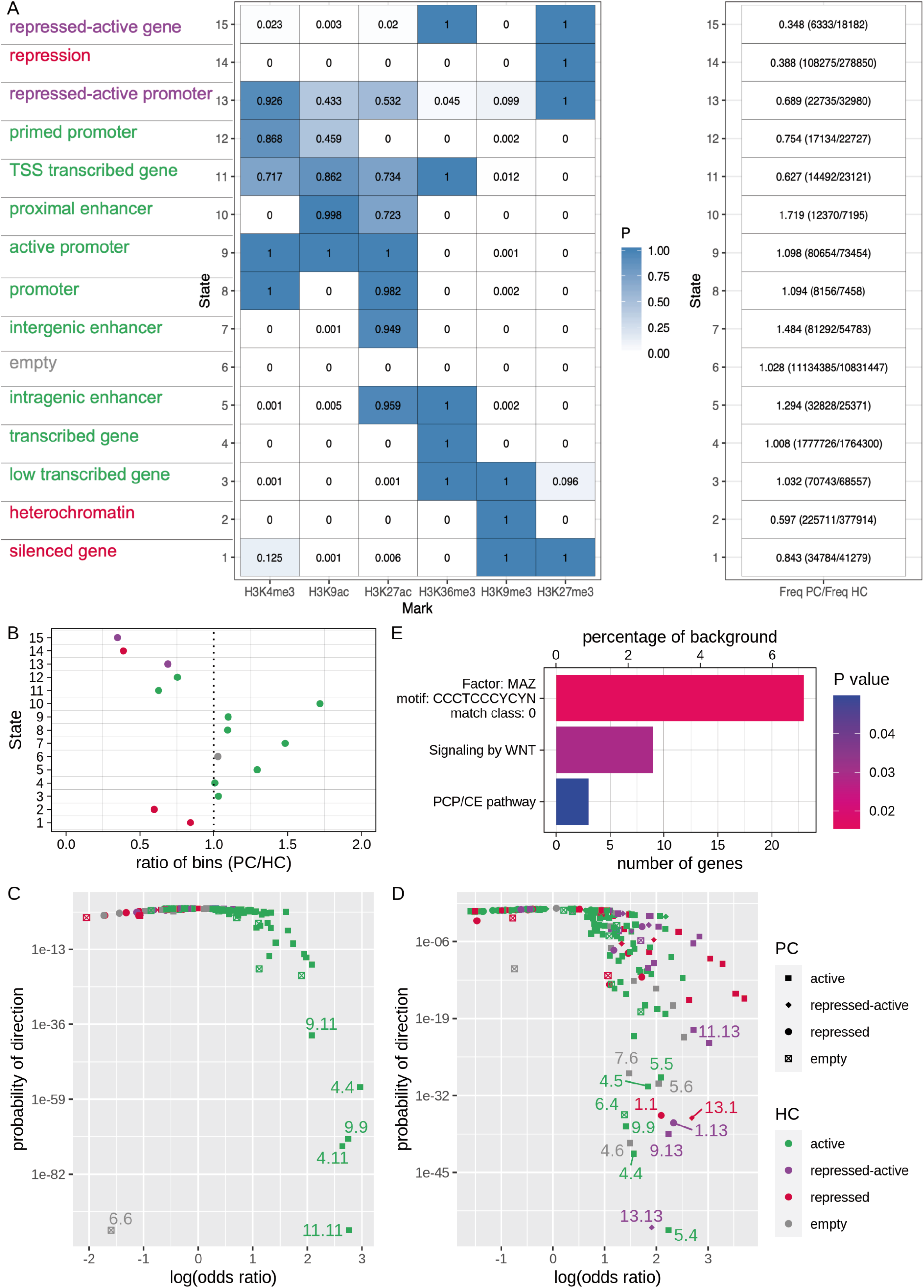
Analysis of ChromHMM states revealed an enrichment of transitions into repressed-active states. (A) Emission probabilities of a 15-state ChromHMM model based on the conserved PC and HC datasets. Leftmost column: functional annotations of states with activating states (green), repressed states (red), repressed-active states (purple) and empty state (grey). Right column: ratio and frequencies of the respective states in PC and HC (in parentheses). (B) Ratio of genomic bins in each ChromHMM state. (C-D) Enrichment or depletion of each ChromHMM state transition in a given set of 265 housekeeping genes (C) and 334 chondrogenesis-associated genes (D) (given in Suppl.Table 2). The log(odds ratio) is shown against the probability of direction (enrichment or depletion). (E) Enrichment analysis of chondrogenic genes showing a transition from an activating promoter state 9 in PC to the repressed-active promoter state 13 in HC.

**Figure 3:**
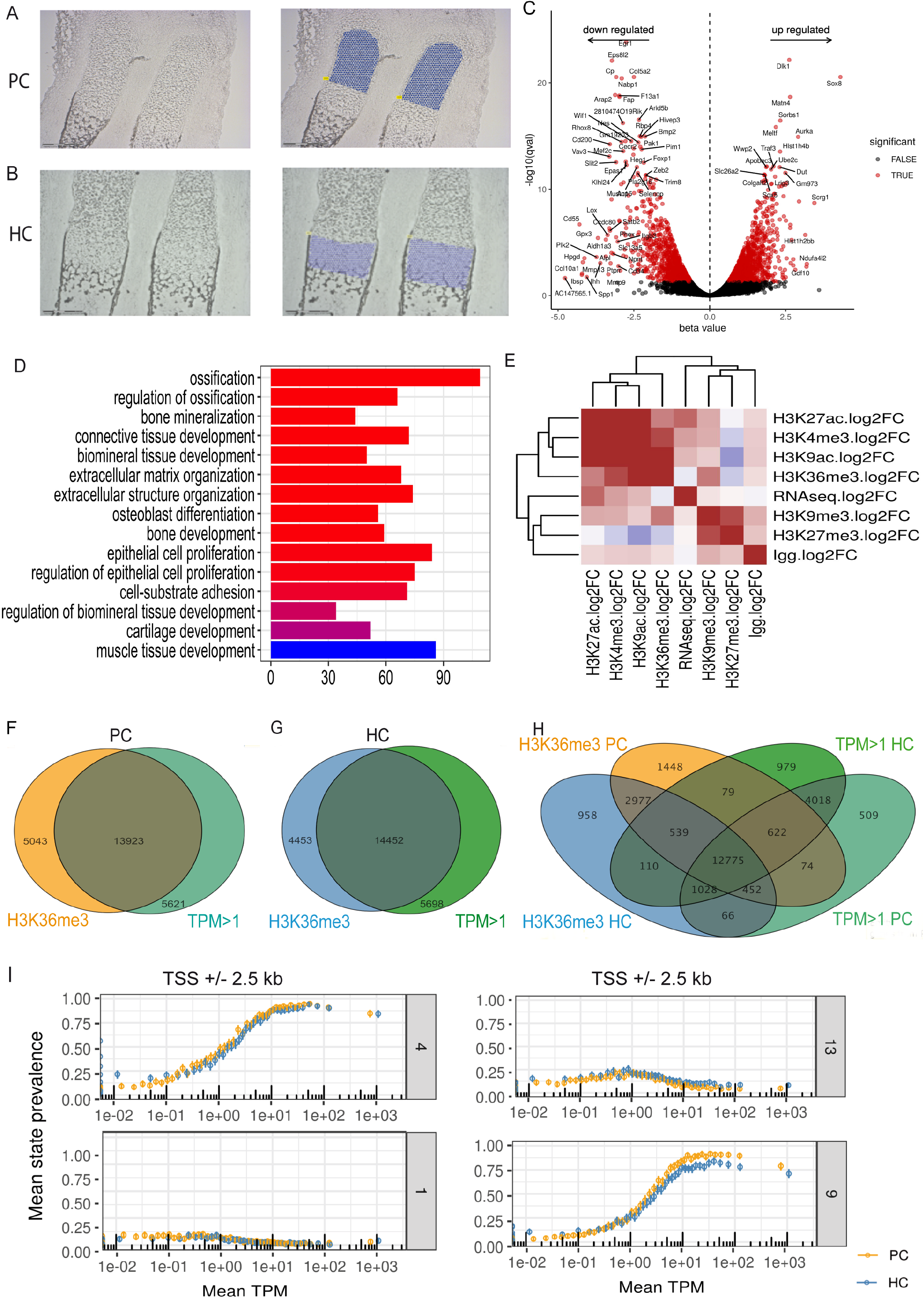
Correlation of differential gene expression and epigenetic modifications. (A-B) Chondrocytes were isolated by laser-microdissection from cryosections of E14.5 (PC, A) or E15.5 (HC, B) forelimbs for RNA expression profiling. Chondrocyte populations were morphologically distinguished. (C) Volcano plot of differential gene expression between PC (beta>0) and HC (beta<0) with names of the 50 genes with highest differences (beta-value) and confidence (−log10 q-value) (D) GO-terms with the highest enrichment of genes differentially expressed between PC and HC. (E) Spearman correlation of differential gene expression and differential occupancy of the indicated histone marks. (F-H) Venn diagrams for expressed genes measured in RNA-seq (green) and genes carrying H3K36me3 in PC (orange) or HC (blue) as marker for transcription in PC (F) and HC (G) and between all four datasets (H). (I) TPM values of genes assigned to a specific ChromHMM state (1, 4, 9, 13) at the promoter region (TSS +/− 2.5 kb) in PC (orange) and HC (blue). Mean state prevalence was plotted against mean log(TPM).

We next asked if hypertrophic differentiation is associated with distinct changes in the epigenetic profile. Comparing the histone states of the two cell populations revealed more states defined by activating marks in PC, while repressive and repressed-active states were more frequent in HC (Fig. 2A-B). This is in good agreement with the observation that genes downregulated in HC gain H3K27me3, while maintaining activating histone marks (Fig. 1 G-H). We thus tested if this gain in repressive states is characteristic for chondrocyte-specific genes by calculating the enrichment of ChromHMM state transitions between PC and HC in a set of 334 chondrogenesis-associated genes and compared it to the enrichment in a set of housekeeping genes (21) (Suppl.Table 2). For the housekeeping genes, transitions between the activating states 4, 9 and 11 showed the highest enrichment (Fig. 2C, Suppl.Fig. 3E). Likewise, transitions between activating (5.4, 4.5), activating and empty (4.6, 6.4, 5.6, 7.6), and the same (1.1, 4.4, 5.5, 9.9 and 13.13) states were enriched, in the set of chondrocyte-specific gene, likely reflecting the similar expression pattern in the closely related cell types (Fig. 2D, Suppl.Fig. 3F). Transitions between state 1 and state 13 also occurred rather frequently in both directions, pointing to a high variability of marks on permanently repressed genes (Suppl.Table 3). Importantly, transitions from the activating promoter states 11 and 9 to the repressed-active state 13 were unidirectionally enriched in the chondrocyte-specific dataset (Fig. 2D). Genes undergoing this transition included several genes known to be downregulated in HC like Sox5, Sox6, Sox9 and Ptch1 (Suppl.Table 3). Enrichment analysis revealed an association to non-canonical Wnt-signaling and Myc/Maz-transcription factors, which have been linked to the switch from PC to HC (Fig. 2E) (22, 23). In summary, the gain of H3K27me3 on promoters still decorated with activating marks seems to be specifically linked to the repression of transcription.

### The gain of H3K27me3 initiates gene repression

To correlate changes in epigenetic states and gene expression, we analyzed the transcriptome of PC and HC isolated by laser microdissection from E14.5 and E15.5 mouse forelimbs with RNA-seq (Fig. 3A-B). Similar read numbers were obtained for all datasets and a high correlation was detected between replicates (Suppl.Fig. 4A-B). Many genes known to be differentially expressed, like *Sox5* and *Matn4* in PC and *Col10a1* and *Mmp9* in HC were found in the respective dataset (Fig. 3C). Furthermore, differentially expressed genes were highly enriched in GO-terms related to bone development and ossification reflecting the chondrogenic cell type (Fig. 3D). Spearman-rank correlation detected a positive correlation between the RNA-seq data and the activating, but not the repressive histone marks (Fig. 3E). On gene level, H3K36me3 occupancy overlapped with transcribed genes identified by RNA-seq in both cell types (Fig. 3F-H). Interestingly, we also found a positive correlation between the differential levels of gene expression and the occupancy of activating promoter and enhancer marks. Furthermore, differences in H3K27me3 negatively correlated with those of the activating marks (Fig. 3E). In contrast, the differential H3K36me3 occupancy did not correlate with changes in gene expression (Fig. 3E) indicating that the presence but not the level of H3K36me3 is linked to gene expression.

**Figure 4:**
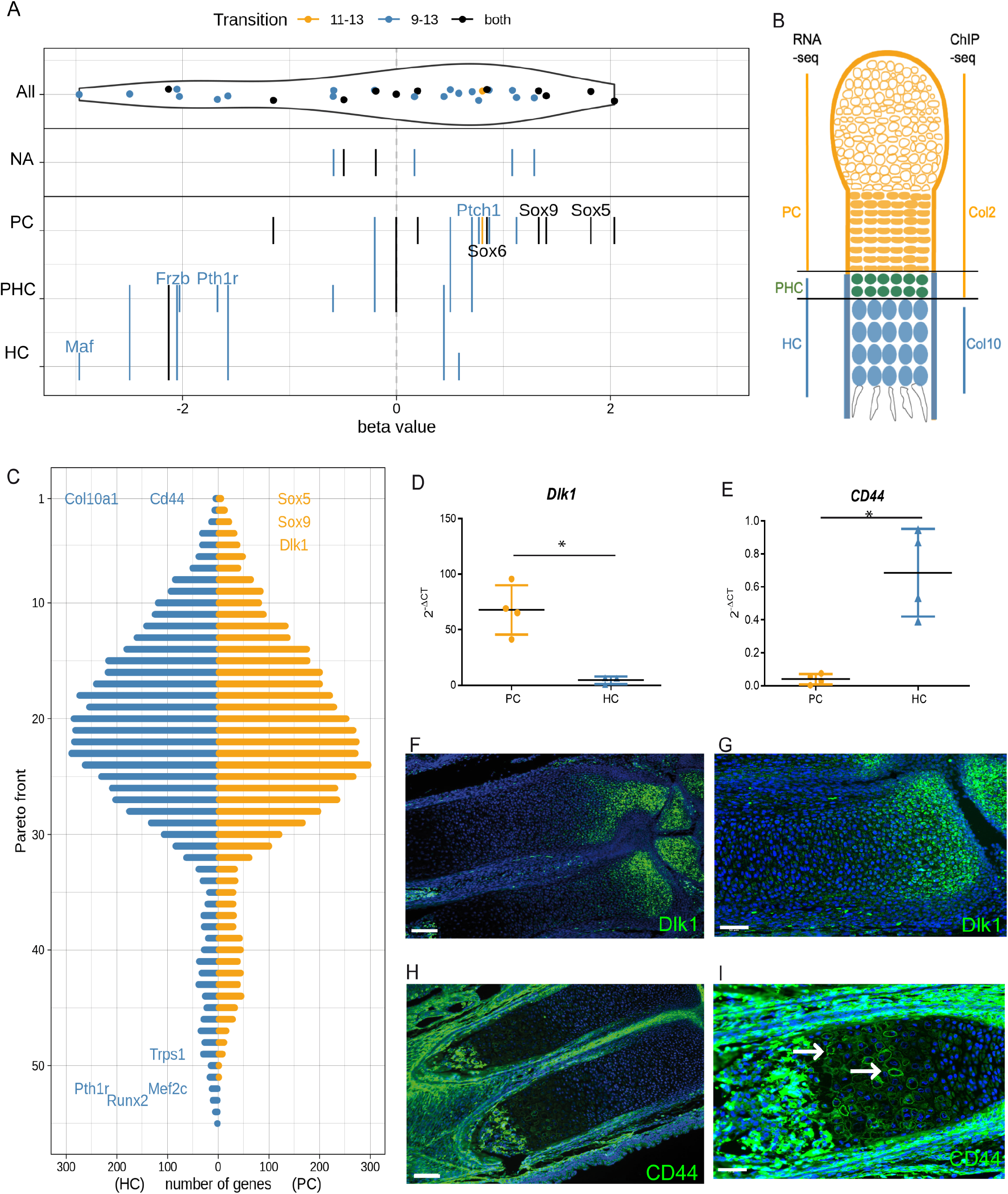
Integration of RNA-seq and ChIP-seq data (A) Chondrocyte-specific genes with the transition from state 9 to state 13 (blue), state 11 to state 13 (orange) or both transitions (black) were sorted by their expression level (beta-value) and assigned to their published expression pattern in PC, PHC and HC or not annotated (NA). Names are given for representative chondrocyte marker genes. (B) Model depicting differences in chondrocyte isolation between the ChIP-seq and the RNA-seq datasets. The methodological difference allowed us to discriminate the pool of prehypertrophic cells (PHC) in each method. (C) Number of genes at a given Pareto front from 1 to 55 with front 1 being the optimal. Genes are color coded according to their differential expression in PC (orange) and HC (blue). Representative genes of the topmost- and lowest-ranking Pareto fronts are given. (D-E) Differential expression of *Dlk1* (D) and *CD44* (E) mRNA was measured by qPCR, detecting higher expression of *Dlk1* in PC (orange) and *CD44* in HC (blue). The −log2^DCT^ is given. n=4 biological replicates, p: 0.0013 (D), p:0.022 (E), error bars show the standard deviation. (F-G) In situ hybridization of *Dlk1* on sections of E16.5 forelimbs detected expression in PC. (H-I) Immunofluorescence staining of CD44 on E16.5 forelimb sections revealed CD44 protein in the cell membrane of HC in the periosteum and the bone marrow. Nuclei were counterstained with DAPI. Scale bars: 100 μm (F, H) 50 μm (G, I).

Analyzing the distribution of ChromHMM states, we found a positive correlation between TPM values and the prevalence of promoter and enhancer states and the transcription-associated state 4 on the promoter (TSS +/−2.5 kb), gene body and enhancer (gene body +/−10 kb) regions, while repressive histone states were enriched on genes with low TPM values. Similarly, the repressed-active state 13 was prevalent on genes with intermediate to low TPM values (Fig. 3I, Suppl.Fig. 4C). Based on the transition analysis we hypothesized that the gain of H3K27me3 associated with the transitions 9.13 and 11.13 leads to the downregulation of PC-specific genes in HC. To test this hypothesis, the differential expression of genes undergoing this transition was compared to the published expression patterns and many of these genes, like *Sox9* and *Sox5*, showed the expected downregulation in HC (Fig. 4A, Suppl.Table 3).

ChromHMM states combine histone modifications at specific genomic locations, but the number of reads detected for each modification is not considered. To quantitatively correlate alterations in epigenetic modifications with differential expression, we integrated the differential level of RNA-seq and promoter-specific ChIP-seq reads (TSS +/−2.5 kb) by Pareto optimization. As the differential H3K36me3 and H3K9me3 occupancy did not correlate with differential gene expression (Fig. 3E) they were excluded from the analysis. The genes were sorted into Pareto fronts based on their positive (for the same direction) or and negative (for H3K27me3) z-scores (Suppl.Table 4). Many genes known to be differentially expressed, like *Sox5*, *Sox9* and *Col10a1,* were found in the topmost-ranking Pareto fronts (Fig. 4C). These fronts also included many yet uncharacterized genes with epigenetic profiles similar to those of *Sox5* and *Sox9,* like *Delta-like kinase 1* (*Dlk1*; Suppl.Fig. 4D) in PC or to *Col10a1, like CD44*, in HC (Suppl.Fig. 4E). The cell type specific expression of these genes was confirmed by qPCR and *in situ* hybridization or immunohistochemistry (Fig. 4 D-I). Together these results strongly support the hypothesis that gene repression is initiated by the gain of H3K27me3 on promoters still carrying a combination of at least 3 activating marks.

Surprisingly, the lowest-ranking Pareto fronts included several genes, like *Runx2*, *Mef2c, Trps1* and *Pth1r,* which are specifically upregulated in cells undergoing the state transition (prehypertrophic chondrocytes, PHC) and are subsequently downregulated in HC (Fig. 4C, Suppl.Table 4). These genes also undergo the transition from an active to a repressed-active state (Fig. 4A, Suppl.Table 3). The repressed-active state in HC corresponds to the downregulation of expression in these cells. Closer inspection of individual Z-scores revealed a negative correlation of differential mRNA expression and activating marks indicating that these cells were included in different cell pools used for RNA-seq (HC) and epigenetic characterization (PC) (Fig. 4B). While supporting the repressed-active state as an indicator of early gene repression the different sampling methods lead to a specific accumulation of transition associated genes in the lowest-ranking Pareto fronts (Fig. 4C) making them a source for new potential regulators of the switch in cell state.

### HC-specific enhancers regulate metabolic pathways

Based on H3K27ac occupancy (24) we defined three states as enhancer states, the proximal enhancer state 10, which included the promoter mark H3K9ac and was mainly found close to the TSS, and the distal enhancer states, state 5 (intragenic enhancer) and state 7 (intergenic enhancer), which were located at distances of up to >500 kb outside the next TSS (Fig. 5A-B, Suppl.Fig. 5A). All enhancer states were more prevalent in PC and correlated positively with the expression of the nearest gene (+/−1 Mb) in both cell types (Fig. 2A-B, Suppl.Fig. 4C). Classifying enhancers as common (22%) PC-specific (51%) and HC-specific (27%) revealed a substantial difference between the cell types (Suppl.Fig. 5B). Comparing the genomic distribution revealed the expected localization close to the TSS for all proximal enhancers, while HC-specific, distal enhancer states are located closer to the TSS (5-50 kb) than PC-specific, distal enhancer states, which are more frequently found at larger distances (>500 kb) (Fig. 5A-B). Enrichment analysis using GREAT assigned proximal enhancer states of both cell types to genes regulating cellular functions like DNA-, RNA- and protein-metabolism (Fig. 5C). The common, distal enhancer states were linked to pathways relevant for both cell types, like skeletal and blood vessel development. Interestingly, PC-specific, distal enhancer states were associated with genes of chondrogenesis-related pathways, while the identified HC-specific, distal enhancer states negatively regulated metabolic processes, likely required for the altered cell volume and ECM remodeling in HC (Fig. 5D).

**Figure 5:**
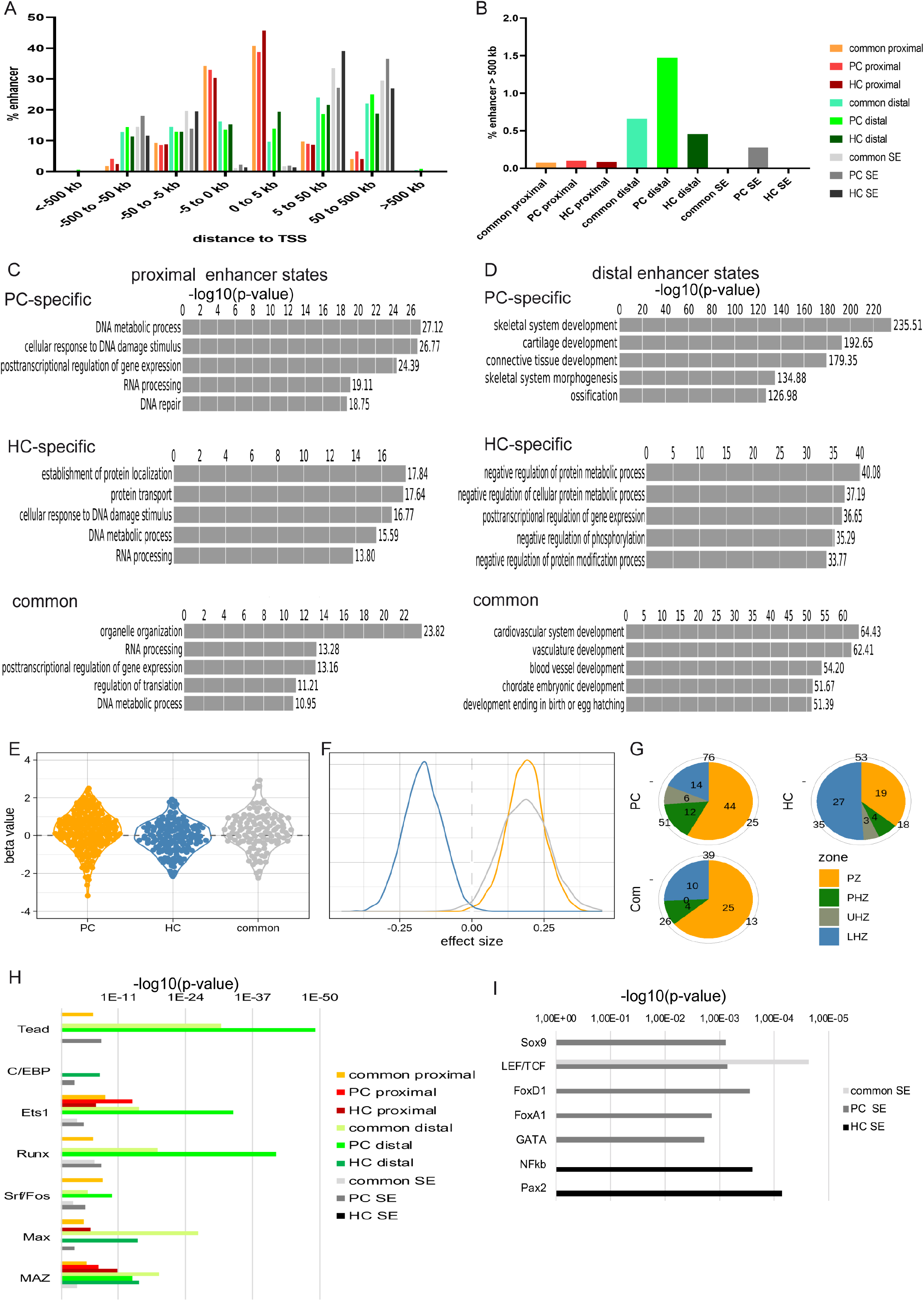
Characterization of enhancers and super-enhancers in PC and HC. (A-B) Distribution of proximal and distal enhancer states and SEs at a given distance from the assigned gene. The enhancer states were separated by their occurrence in either cell type-specific (PC or HC) or common enhancers. (C-D) Enrichment analysis of genes annotated to either proximal (C) or distal (D) enhancers showing the five GO-terms with the highest enrichment for PC-specific, HC-specific or common enhancers. Enhancers were assigned to the nearest gene within 1000 kb using the GREAT algorithm. (E-F) Differential gene expression (E) and effect size of the correlation (F) between genes regulated by PC-specific (orange), HC-specific (blue) and common (grey) SEs compared to differential expression of all genes. (G) Pie chart of the distribution of SE-carrying genes in PC, HC or common in both in relation to their published expression pattern in the proliferating zone (PZ), prehypertrophic zone (PHZ), upper hypertrophic zones (UHZ) and lower hypertrophic zones (LHZ). Information about gene expression was derived from Tan et al. (26). The total number of enhancers in the specified zone and cell type is given. (H) Enrichment (−log10 p-value) of predicted transcription factor binding sites found in the different enhancer types. (I) Enrichment (−log10 p-value) of the predicted transcription factor binding sites found in SEs.

Super-enhancers (SEs), defined as clusters of H3K27ac peaks (25), are important regulators of differentiation. We detected 360 SEs specifically in PC, 205 in HC and 158 shared by both cell types. Assignment of SEs to the nearest gene identified tissue-specific SE associated with regulators of chondrogenesis, like *Sox5*, *Sox9* and *Col10a1*, and many new potential regulators (Suppl.Table 5). We identified 30 genes, including *Tcf4* and *Smad7*, that carried cell type-specific SEs at different genomic locations indicating different activating mechanisms in the respective cell type (Suppl.Table 5). Enrichment analysis assigned similar biological functions to SEs as to distal enhancer states, chondrogenic differentiation for PC-specific SEs and metabolism for HC-specific SEs (Suppl.Fig. 5C). On average, genes carrying a cell type-specific SE were expressed at significantly higher level in the respective cell population (Fig. 5E-F, Suppl.Fig. 5E) with several PHC-specific genes being associated with PC-specific SEs (Suppl.Table 5). Comparing the relationship of SEs to independently obtained expression data (26) confirmed the association of PC-specific SEs to genes expressed in PC and PHC, while HC-specific SEs were mainly associated with genes found in the lower hypertrophic region (Fig. 5G).

To gain insight into the transcriptional control of chondrocyte differentiation, we screened the enhancer states for predicted transcription factor binding motifs. We found binding motifs for Maz, Max, Srf/Fos, Runx, Ets1, C/EBP and Tead, general regulators of differentiation, differentially enriched in both cell types (Fig. 5H). In accordance with their role as main regulators of differentiation, SEs included additional binding sites for chondrogenesis-related transcription factors like Sox9, LEF/TCF, FoxD1, FoxA1 and GATA in PC and NFkB and Pax2 in HC (Fig. 5I).

As developmentally relevant transcription factors frequently bind to less conserved binding motifs (16, 27), we compared Sox9 ChIP-seq data (16) to our enhancer states. About 16% of the proximal enhancer states overlapped with Sox9 peaks. Of the distal enhancer states, 31% of the common and 22% of the PC-specific states colocalized with Sox9 peaks, while only 13% of the HC-specific states showed an overlap (Fig. 6A). Furthermore, nearly all SEs colocalized with Sox9 peaks in both cell types (Fig. 6A, Fig. 6B) supporting the hypothesis that establishing the Sox9 dependent chondrogenic fate is the main biological function of PC, while the differentiation of HC is dependent on the activation of metabolic pathways.

**Figure 6:**
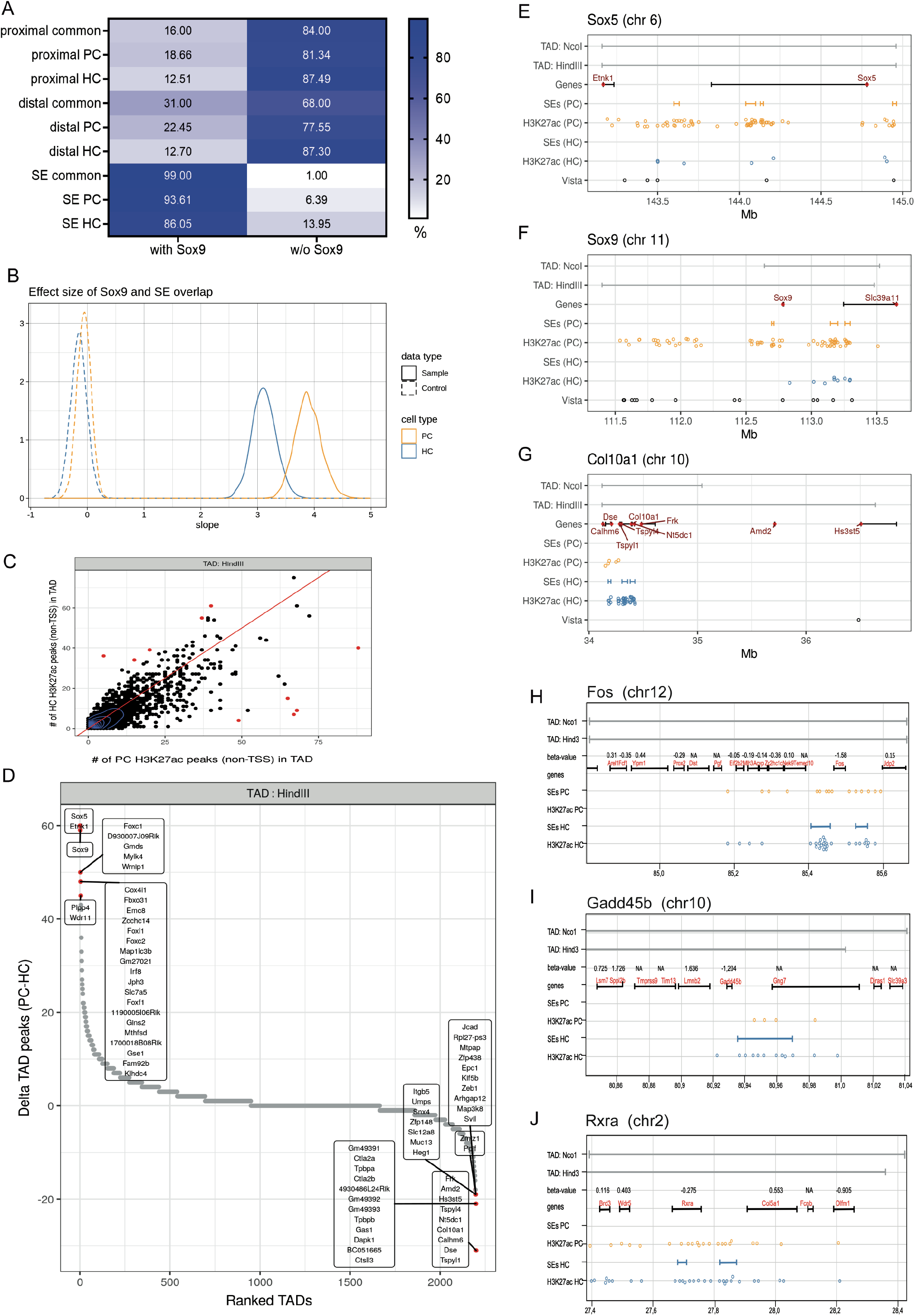
Differential H3K27ac occupancy within a TAD correlates with cell-type specific gene expression. (A) Colocalization of Sox9 peaks with different enhancer states and SEs in PC and HC. (B) Effect size of the Sox9 enrichment at SEs in both cell types compared to randomly chosen genomic regions of similar size. (C) Enrichment of enhancer peaks in TADs. The number of H3K27ac peaks outside the TSS was compared in PC and HC. (D) TADs were ranked according to their differential occupancy with H3K27ac (deltaTAD peaks). (E-I) H3K27ac peaks (circles) and SEs (bars) detected in PC (orange) and HC (blue) within TADs (grey bars) containing Sox5 (E), Sox9 (F) Col10a1 (G), Fos (H) Gadd45b (I) and Rxra (J). Distances are given in Mb.

### Cell state transitions are associated with a rapid change in enhancer usage

Enhancers have been predicted to act on genes within the same topologically associating domain (TAD) (28) with the number of H3K27ac peaks within a TAD correlating with the strength and the robustness of gene expression (25, 29). To gain further insight into the importance of enhancer coverage on gene expression, we calculated the number of H3K27ac peaks within a TAD (30) and ranked TADs according to the ratio of H3K27ac peaks between PC and HC (Fig. 6C-D, Suppl.Fig. 5F-G, Suppl.Table 6). The TADs with the greatest differential number of H3K27ac peaks contained the genes *Sox5*, *Sox9* and *Col10a1*. Other TADs with high differences in enhancer marks contained transcriptional regulators like Fox-family proteins in PC and Tead1 in HC (Fig. 6D, Suppl.Fig. 5G, Suppl.Table 6) marking them as potentially important regulators of chondrocyte differentiation. In agreement with the decline in *Sox9* and *Sox5* expression during hypertrophy, the clustering of enhancer peaks observed in PC was lost in HC, but few enhancer peaks were still maintained (Fig. 6E-F). In contrast, the TAD covering the *Col10a1* gene contained few H3K27ac peaks in PC, while their number was increased, and several clusters were identified in HC (Fig. 6G). While several Sox5 and Sox9 associated enhancers matched conserved enhancers of the Vista database none of these HC-specific enhancers were included (Fig. 6E-G), indicating that less conserved mechanisms regulate hypertrophy. To test the correlation of peak accumulation and expression changes, we inspected the genomic location of genes like *Fos, Gadd45b* and *Rxra*, which were upregulated and linked to SEs in HC (Fig. 6H-J). As observed for *Col10a1*, the genomic regions were pre-marked with H3K27ac peaks in PC, pointing to a priming of enhancer clusters prior to gene expression and a fast accumulation of additional marks with the onset of gene expression.

## Discussion

Previous studies of chromatin remodeling focused mainly on the *in vitro* differentiation of embryonic and mesenchymal stem cells into different cell lineages (1, 2), but insight into the epigenetic regulation of distinct differentiation steps within one cell type are less well understood. In this study, we have, for the first time, investigated changes in the epigenetic profile linked to a specific differentiation step, the terminal differentiation of proliferating into hypertrophic chondrocytes, and have correlated these to differential gene expression.

Analyzing a set of activating and repressive histone marks, we generated an unbiased map of epigenetic modifications. Interestingly for H3K4me3, H3K9ac, and H3K27ac not only the coverage but also the differential level of modification correlated with differences in gene expression identifying them as predictors for changes in the transcriptional activity. H3K27me3 negatively correlated with the activating promoter marks, indicating that its differential occupancy also affects gene expression. Combining histone marks into states by ChromHMM identified two states that combine the repressive mark H3K27me3 with either activating promoter marks (state 13) or H3K36me3 (state 15). Analyzing the frequency of state transitions between PC and HC in a set of chondrogenesis-related genes revealed a specific enrichment of transitions from activating states 9 and 11 into state 13 on genes like *Ptch1, Sox5, Sox6* and *Sox9*, which are downregulated during hypertrophy. Furthermore, integrating differential expression and epigenetic modifications identified many genes known to be downregulated in HC in the topmost-ranking Pareto fronts, supporting the hypothesis that the addition of H3K27me3 on regions still marked as active and not the removal of activating marks is the repression-inducing event.

A gain of H3K27me3 at H3K4me3-decorated promoters has recently been described for neuronal genes downregulated in the process of neural differentiation (31). The presence of other activating marks, which were included in our data, has not been investigated. It is likely that during differentiation neuronal progenitor cells, like chondrocytes, add H3K27me3 to promoters marked as active by other modifications besides H3K4me3. Sodersten et al., 2018 (31) also detected an enrichment of the combination H3K27me3/H3K9me3 (corresponding to our state 1) on genes that are permanently silenced in differentiated neurons. In our dataset we found an enrichment of the transitions 1.13 and 13.1 (Fig. 2D, Suppl.Table 3). These transitions were mostly associated with early limb patterning genes, which are no longer expressed in either chondrocyte type, and likely reflect the transition into permanent repression with rare leftovers of activating marks. Surprisingly genes downregulated between PC and HC were not enriched in the transition into the repressed state 1. This difference to the neuronal data likely reflects the longevity of neurons, while HCs have a life span of a few days before they undergo apoptosis or transdifferentiate into osteoblasts (32, 33). They might thus not have sufficient time to convert the epigenetic decoration from repressed-active into permanently silenced. In neuronal cells, genes repressed by H3K4me3/H3K27me3 (31) are preferably reactivated during neuronal stress. Recent studies demonstrated that hypertrophic chondrocytes undergoing ER stress, can re-enter the proliferating state (34, 35). One might thus speculate that the incomplete loss of activating marks facilitates this redifferentiation during pathological conditions.

While the repression of genes is linked to the repressed-active chromatin modification, we did not detect an enrichment from a bivalent to an activating state in genes upregulated in PC as it has been observed for lineage-specific genes in ES cells (36). Inspection of CD44, which we identified as a new maker for HC, did not show repressive marks at the promoter region in PC. The bivalent modification preparing for expression might thus be preferentially found in ES cells (37), while at later developmental stages, lineage-specific genes have lost the pluripotency associated label. In summary, while we did not find an enrichment of repressed-active modifications on genes prone to be upregulated in HC, we gained strong evidence that the initiation of gene repression is linked to the addition of the repressive mark H3K27me3 to regions still marked as active, identifying the addition of this mark rather than the removal of activating marks as the repression inducing event. Long term repression might subsequently require the addition of H3K9me3 as indicated by the neuronal data.

To gain further insight into the cell state specific control of gene expression, we identified enhancers based on H3K27ac occupancy. Although both, PC and HC, belong to the chondrogenic lineage, the identified enhancers are quite different, with only about 30% common to both cell types. GO analyses gave insight into the chondrocyte specific function of enhancer associated genes. As expected, PC-specific enhancers were enriched in chondrogenesis related pathways establishing the specific cell lineages. Intriguingly, continuous activation of lineage defining genes is less important in HC, after the chondrogenic lineage has been established. The loss of lineage-specific enhancers might be indicative for the terminal differentiation of a cell type. In contrast to PC, HC specific enhancers related mainly to metabolism associated genes. Little is known about the mechanisms by which HC acquire their size and specific ECM. Increased metabolic activity (38, 39), Bmp dependent mTor (40, 41) and IGF1 signaling (42, 43) have been implicated in regulating hypertrophy before and our data strongly emphasize the importance of metabolism for the generation of hypertrophy.

The association of enhancers to the regulated gene is hampered by their function over long distances, which may also include other genes (18, 44), (45). It is thus difficult to predict if specific enhancers control different genes or if different enhancers regulate the same gene at distinct differentiation stages. Inspection of SEs indicates that both may be the case. Differential expressed genes, like Sox9 and Col10, were associated with cell type specific SEs, while genes like Tcf4 and Smad7, were assigned to different SEs in PC and HC. More direct investigations of enhancer usage by e.g. unbiased chromatin conformation capture (46) will be required to gain a detailed understanding of their cell type specific use, interaction and function.

Several studies have linked the number of enhancers to the level and robustness of gene expression (47). To analyze differences in enhancer usage, we ranked TADs by their differential H3K27ac abundance. The highest-ranking TADs contained cell type-specific genes like Sox9, Sox5 and Col10, while genes like Smad7 and Tcf4, regulated by distinct SEs in either cell type, were found in average-ranked TADs. Inspection of single TADs confirmed that the downregulation of genes like Sox9 and Sox5 is linked to a rapid loss of enhancers in HC. In contrast, upregulation of genes like Col10 and Gadd45b in HC is accompanied by a low enhancer coverage in PC followed by a dramatic increase in HC. While the association of enhancers to genes was only possible for selected TADs, correlating expression differences of single genes to enhancer usage inside a TAD in more detailed studies will provide valuable insight into the control of gene expression (48, 49). Furthermore, recent studies demonstrated that TADs are not static units, but are remodeled dependent on transcription (28). Establishing chondrocyte-specific TADs might thus refine the interpretation of enhancer association and usage. In summary, while differences in enhancer usage have been reported for genes expressed in different cell lineages (50), our data provide first insight into the dynamics of enhancer usage in sequential differentiation stages.

## Material and Methods

### Animal models

Collagen 2a1-Cre (Tg(Col2a1-cre)1Star (51)), Collagen 10-Cre mice (Tg(Col10a1-cre)1427Vdm, (52) and ROSA26-YFP (Yellow fluorescent protein) reporter (R26R-YFP, ((Gt(ROSA)26Sortm1(EYFP)Cos, (53) mice were maintained on a C57Bl/6J genetic background and crossed to generate time pregnancies of Col2-Cre;R26RYFP and Col10-Cre;R26R-YFP mice. Mouse husbandry was approved by the city of Essen (Az.32-2-11-80-71/203) in accordance with §11 (1) 1a of the “Tierschutzgesetz” and approved by the animal welfare committee of the University Duisburg-Essen. Embryos were decapitated, freed from skin, muscle and internal organs, dissociated with 0.6 U/ML Collagenase (Serva) and 0.25 Trypsin (Thermo), filtered through a 70 µm cell strainer (Falcon) and fixed with 1% formaldehyde (Applichem) prior to cell sorting to eliminate potential, FACS-induced stress response and corresponding alterations in chromatin structure.

### Cell sorting and Chromatin immunoprecipitation with next generation sequencing (ChIP-seq)

PC and HC were isolated from E13.5 Col2-Cre;R26R-YFP and E15.5 Col10-Cre;R26R-YFP embryos, respectively, by FACS of YFP positive cells using a FACSAria II cell sorter (BD Bioscience) with a 70 μm nozzle. 3,5×10^5^ cells were used for each ChIP. Chromatin was sonicated for 30 min using a Biorupter (Diagenode) and isolated using the True MicroChip Kit (Diagenode). Antibodies for ChIP were obtained from Diagenode (H3K4me3, H3K36me3, H3K9me3me, H3K27me3, IgG) or Abcam (H3K9ac, H3K27ac). Library preparation was performed with the MicroPlex Library preparation Kit V2 (Diagenode) and 61 bp reads were sequenced on an Illumina Hiseq 2500 (Illumina) at a depth of 20-30 million reads per experiment. Biological triplicates were performed in triplicate for PC, each replicate representing cells of approximately 10 embryos, and in duplicate for HC including cells of approximately 50 embryos per replicate.

### ChIP-seq data processing

Read quality of the raw sequences was verified with fastqc (version 0.11.9, (Andrews, 2010). All reads were trimmed to a length of 60 bases and those with a mean Phread score below 20 were discarded using prinseqlite (version 0.20.4, (Schmieder and Edwards, 2011)). The remaining reads were mapped to the mm10 build (UCSC mm10, Dec. 2011) of the mouse reference genome with bwa (version 0.7.15, (Li and Durbin, 2009)) using the default setting, except the maximum edit distance, which was set to 4. Unmapped reads and reads with a mapping quality below 30 were removed with samtools (version 1.4/1.5, (Li and Durbin, 2009). Duplicate reads were filtered with Picard tools (version 2.9, http://broadinstitute.github.io/picard/). Mapping quality statistics were computed using qualimap (version 2.1 (Kharchenko et al., 2008; Okonechnikov et al., 2016)). To assess the library complexity, the Non-Redundant Fraction (NRF) and PCR Bottlenecking Coefficients (PBC, https://github.com/imbforge/encodeChIPqc/blob/master/R/PBC.R) were calculated with functions implemented in R (version 3.6.1). Relative Strand Correlation (RSC) and Normalized Strand Correlation (NSC), which indicate efficiency of the ChIP, were determined using Phantompeakqualtool (54, 55), (https://github.com/kundajelab/phantompeakqualtools). Based on the FRiP (fraction of reads in peaks) parameter (55) the fraction of reads in enriched regions (FRIER,https://github.com/BioinformaticsBiophysicsUDE/ChIPSeqQualityMetrics/blob/master/FRIER.R) was calculated. Briefly, the coverage fold-change over the global background in 3000 bp windows was computed in an interval from 0.1 to 4.0 using R-packages csaw and GenomicAlignment (Lawrence et al., 2013; Lun and Smyth, 2016). The concordance between biological replicates was computed as Pearson and Spearman correlation (https://github.com/BioinformaticsBiophysicsUDE/ChIPSeqQualityMetrics/blob/master/Spearman_Pearson_Correlation.R). For the mapped and filtered reads the coverage was binned (5 kb bins) and the correlation was calculated with R. Quality parameters were set to match the criteria defined by the Encode consortium. The code is available at: https://github.com/BioinformaticsBiophysicsUDE/ChIPSeqProcessingOld.

Peaks were identified with hiddenDomains (version 3.0 (19)). As a control dataset all IgG data for one cell type were merged. The bin size was adjusted to 200 bp (narrow domains for H3K4me3, H3K9ac, H3K27ac) or 800 bp (broad domains for H3K36me3, H3K9me3, H3K27me3). The parameter for the maximum read count was decreased to 30 and the posterior probability threshold set to 0.6. Using a sliding window approach, peaks separated by one or two bins, for broad and narrow domains, respectively, were merged. Consensus peaks were obtained by taking the intersection of the biological replicates. Comparability between the different replicate numbers was tested by calculating the peak number and coverage for subsets of two PC replicates. Code available at: https://github.com/BioinformaticsBiophysicsUDE/ChIPSeqPeakCalling. The Integrative Genomics Viewer (IGV, http://software.broadinstitute.org/software/igv (56)) was used to visualize histone marks and peaks at selected genomic locations.

### Defining epigenetic states with ChromHMM

The peaks generated by hiddenDomains were used as input for ChromHMM (version 1.20 **(57)**). Each chromosome was segmented into non-overlapping bins of 200 bp. For each histone mark a bin was set to 1 if it partially or completely overlapped with a peak, and to 0 in case of no overlap generating a separate track of bins for each histone mark. This data was used to train multiple ChromHMM models with increasing number of states. Based on their log-likelihoods and the biological interpretation the 15-state ChromHMM model was selected. To assess the robustness of the model the duplicate subsets of the PC an HC datasets were evaluated. Additionally, the Encode mouse reference epigenomes for limb (ENCSR283NCE, ENCSR589KOM, ENCSR424HQH, ENCSR705YBY, ENCSR486MYL) and forebrain (ENCSR415TUB, ENCSR296YDZ, ENCSR911FIM, ENCSR894URD, ENCSR438IXZ) from developmental stages E11.5-E15.5 were included. Code available at: https://github.com/snaketron/SourceChromHmm/code.zip

### Genomic enrichment of ChromHMM states

The enrichment of ChromHMM states was quantified for in ten functional annotations: RefSeq transcription start site (TSS), transcription end site (TES), regions within 2 kb around the TSS, exons, genes, CpG islands, VISTA (www.enhancer.lbl.gov; accessed 2018), Fantom enhancers (https://fantom.gsc.riken.jp; version 2015) and Zinc finger genes (ENSEMBL annotation for the keyword ‘zinc finger protein’), all enhancers” describes Fantom enhancers present in at least one tissue, “forelimb” are Fantom enhancers detected in forelimb only. Enhancer coordinates were converted to mm10 with liftover (https://genome.ucsc.edu/cgi-bin/hgLiftOver). For each ChromHMM state the bin was set to 1 if it partially or completely overlapped with the genomic region assigned to the functional annotations. In case of no overlap, the bin was set to 0. The enrichment of a state at a given annotation compared to the background was computed as E_ij_ = (F_ij_/F_i_)/(F_j_/F); where i and j index different ChromHMM states and functional annotations; F represents the frequency of bins **(58)**

### Enrichment analysis of ChromHMM state transitions

For ChromHMM state transition analysis of PC to HC, genes of the Gene Ontology (GO) terms, chondrocyte differentiation (GO:0002062), limb development (GO:0060173) and cartilage development (GO:0051216) (334 genes, Suppl.Table S2) and, as a control, a group housekeeping genes (265 genes (21) Suppl.Table S2) was selected. For these genes, the occurrence of a given ChomHMM transition was compared to the occurrence in the full set of genes. From this data a binomial regression model was fitted in a Bayesian framework (brms (2.10.0) to determine an enrichment or depletion of a transition in a given set of genes with default priors (59) and rstan (2.19.2) (Stan Development Team. 2018. *RStan: the R interface to Stan*. R package version 2.17.3. http://mc-stan.org).

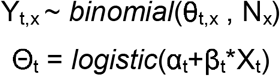

The number of genes with transition t and the condition x (part of the background (x = 0) or the selected group (x =1)) si Y_t,x_. It depends on the number of genes in condition x (N_x_) and the probability θ_t,x_ that a given transition *t* occurs in a gene which is annotated to one of the two conditions x. The influence of the condition on θ is expressed by the effect size β, which is the log odds ratio. The uncertainty that an enrichment (positive log odds ratio) or a depletion (negative log odds ratio) occurs is represented by the probability that β is either negative or positive, respectively. The accuracy of the model was tested by a posterior predictive check and the ability rteocover simulated data. Code available at: https://github.com/keksuntdasteoT/SransitionEnrichment.

### Enhancer analysis

The number of H3K27ac peaks outside the promoter regions (TSS +/−2.5 kb)(60) within a TAD was calculated for PC and HC. The ratio (PC/HC) peaks was used to rank the TADs based on the differential H3K27ac peak number. Information of TADs from embryonic stem cells was obtained from GEO (GSE35156, (30). Pairs of H3K27ac peaks were incrementally clustered together if their distance was lower than 12.5 kb. Clusters with at least five H3K27ac peaks were annotated as SEs (25, 29).Enrichment analysis of enhancer states and SEs was performed using GREAT (version 3.0.0, http://great.stanford.edu/public/html, (61)). Enhancers were assigned to the nearest gene within 1 Mb. Enrichment is given as negative log 10 of the binomial probability parameters.

To quantify the effect of SE on differential gene expression, the β values of the nearest gene was fitted to a linear model (brms (2.10.0) with default priors (59) and rstan (2.19.2)) (Stan Development Team. 2018. *RStan: the R interface to Stan*. R package version 2.17.3. http://mc-stan.org).

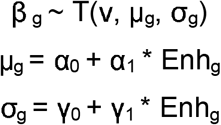

It models the dfiferential gene expression β for a given gene g, by a student-T distribution, defined by ν, μ and σ (degrees of freedom, mean and standard deviation, respectively). The slope α_1_ can be interpreted as the effect the presence of a SE (Enh = 1), has on the differential gene expression compared to its absence (Enh = 0). The accuracy of the model was tested by a posterior predictive check. Code available at: https://github.com/keksundso/SEGeneExpression.

Sox9 peaks (GEO, GSM1692996 (16)) within the same genomic region (200 bp) of enhancer states or SEs were defined as co-localization using StrandNGS (StrandLifeScience, version 2.9). To analyse the enrichment of Sox9 peaks in SEs, the overlap (Y) was compared to that of random shuffled genomic sequences of the same length (bedtools 2.26.0 (62)). As an additional control, the Sox9 peaks where shuffled and the process was repeated. With this data, the following binomial regression model was fitted in a Bayesian framework (brms (2.10.0) with default priors (59) and rstan (2.19.2)) (Stan Development Team. 2018. RStan: the R interface to Stan. R package version 2.17.3. http://mc-stan.org).

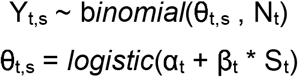

θ_t,s_ is the probability for each type *t* (PC and HC, each original Sox9 peaks and shuffled peaks) that a SE (s = 1) or SE sized peak (s = 0) has a matching Sox9 peak. N is the number of SE peaks and the slope β is the log odds ratio of SE peaks (S = 1) overlapping with Sox9 compared to randomly located genomic regions (S = 0) overlapping with Sox9. The accuracy of the model was tested by a posterior predictive check. Code available at: https://github.com/keksundso/SE_Sox9.

### Transcriptome analysis

Chondrocytes were morphologically distinguished in 10 μm cryosections of E14.5 (PC) and E15.5 (HC) embryonic forelimbs and isolated by laser-microdissection with the PALM Microbeam System (Zeiss MicroImaging). Isolation of RNA from fixed chondrocytes used for the FACS isolation provided insufficient amounts of RNA, while laser-microdissection yielded high quality RNA for sequencing. RNA was prepared with the RNAeasy micro kit (Qiagen). RNA integrity was measured with the RNA 600 pico Kit on a Bioanalyser (Agilent), 10 ng RNA with a RIN>6 was used for library preparation with the NuGen Ovation FFPE library system kit and mouse lnDA-C primers. Sequencing (RNA-seq) was performed on an llumina HiSeq2500 with 101 cycles (PE mode). The quality of the fastq data was examined with FastQC. Trimmomatic-0.36 (63) was used to remove adapters and reads with a quality score below 20. The reads were then pseudo-aligned to mm10 and quantified using Kallisto, version 0.43.1, (64). Expression levels were calculated as transcripts per million (TPM). Transcripts with an expression value below 1 TPM were removed. Differential expression between PC and HC was analysed with Sleuth (65) using the Wald test. Significance was defined as q-value < 0.05, with Benjamini-Hochberg-adjusted false discovery rate to correct for multiple comparisons. Code available at: https://github.com/yingstat/ScriptsforPaperChondrocytes.

GO enrichment analyses were carried out using clusterProfiler (66). Significant enrichment was determined based on q-values as before. Top-enriched GO terms were visualized with clusterProfiler. For the qPCR, cDNA was synthesized with the Maxima First Strand cDNA Synthesis Kit (Thermo Scientific) using 1 ng RNA. qPCR was performed on the CFX384Touch Real-Time PCR detection system (Bio-Rad) with the my-Budget 5x EvaGreen QPCR-Mix II (BioBudget) using primers listed in table 1. Expression data was normalized to Beta-2 microglobulin (B2M), differential expression of four biological replicates was calculated as −2ΔCT. Graphs were plotted with GraphPad Prism (vers.8.4.1). Statistical analysis tested distribution using Shapiro-Wilks test and unpaired, two-tailed Students t-test for significance. Error bars mark the standard deviation between replicates.

**Table1:**
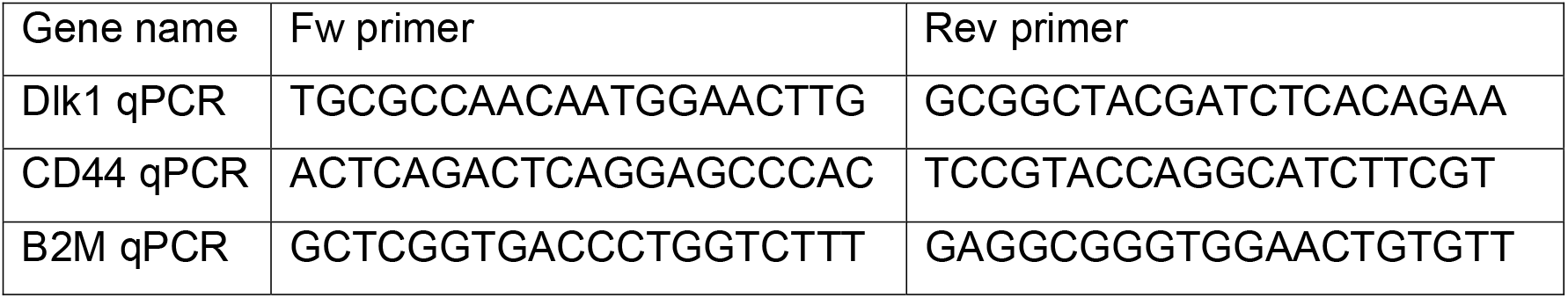
Primers used for qPCR.

### In situ hybridization and immunofluorescence staining

Probes for *in situ* hybridization, were cloned in pCR4 using the TOPO-TA cloning kit (Thermo Fischer Scientific) using the primers Dlk1fw: CCTGGCTGTGTCAATGGAGT, Dlk1rev: GGTGAGGAGAGGGGTACTC (Metabion). RNA antisense probes were labelled with Digoxigenin-11-UTP (Roche). Hybridization was performed as described previously (67). Immunofluorescence staining was performed with a rat-CD44 antibody (Invitrogen) as described previously (68). Fluorescence pictures were taken on a Zeiss Axio Observer7 microscope with an AxioCam 506 mono CCD camera and Zen 2.3 software (Zeiss).

### Correlation between gene expression and ChromHMM state prevalence

The genomic regions (gene: transcribed region, ENSEMBL), TSS (+/−2.5 kb) and enhancer (transcribed region +/−10 kb) of a given gene were described by a binary vector with 15 elements, with each element of the vector representing the prevalence of a particular ChromHMM state, with an element set to 1 if the region overlaps a bin of that ChromHMM state, and 0 otherwise. To associate ChromHMM state prevalence with gene expression levels, the TPM values of each gene were sorted by increasing TPM-values and ranked accordingly. The TPM ranks were split into non-overlapping sets of 500 ranks (genes). For each gene set the mean prevalence of the different ChromHMM states and the mean TPM value was computed using bootstrapping (R-package boot version 1.3-23, number of bootstrap replicates R = 10,000). The 95% highest density intervals (HDIs) of the mean state prevalence is given.

### Integrative analysis of ChIP-seq and RNA-seq data

The integrative analysis was done with R package intePareto (https://cran.r-project.org/web/packages/intePareto/index.html). Counts were estimated from the raw RNA-seq data (FASTQ format) with Kallisto (version 0.43 (64)). The raw ChIP-seq data in FASTQ format were aligned to mm10 with BWA (0.7.15, (69) and sorted with Samtools (version 1.5, (70). To define the epigenetic signal for the number of ChIP-seq reads of each mark falling into the promoter region (TSS +/−2.5 kb) were calculated. Matching of RNA-seq and ChIP-seq data on gene level was performed with “highest” strategy implemented in intePareto. The fold-change was calculated for each histone mark with DESeq2 (version 1.24.0, (71)), and assigned a z-score, reflecting the differential coverage between PC and HC in relation to the fold-change in RNA-seq. Genes were ranked by Pareto front in the space defined by all z-scores. Code available at: https://github.com/yingstat/ScriptsforPaperChondrocytes.

## Supporting information

Supplementay Figurese

## Author Contributions

M.W and A.V. designed the experiments. A.T. and M.W. performed the experiments. A.L. and S.K. analyzed the ChIP-seq data. Y.C. and C.N. analyzed the RNA-seq data. A.L., S.K., Y.C. and C.N. developed the code and provided figures. M.W., C.N. and A.V. interpreted the data. A.M.W provided flow cytometric support. D.H supervised the bioinformatic aspects of the project. A.V. supervised the project. The manuscript was written by M.W., C.N. and A.V. with input from all authors. All authors discussed the results and commented on the paper.

## Acknowledgments

The authors would like to thank Sabine Schneider for excellent technical assistance and critical reading of the manuscript and Dr. Klein-Hitpass for sequencing data. This work was supported by the DFG research unit ExCarBon (FOR 2407; Vo620/14) and the DFG graduate program GRK 1431 to AV.

## Supplementary figure legend

Supplementary figure 1: Quality control of the generated ChIP-seq datasets.

(A) Expected localization of histone marks analyzed relative to the transcription start site (TSS). (B) Principal component analysis of peaks revealed clustering according to the respective histones marks, color scheme as in A, IgG (gray). (C) Spearman correlation between RNA-seq and ChIP-seq data revealed a high correlation between the activating marks and gene expression. The repressive marks positively correlated with each other and with H3K36me3 and negatively correlated with the activating promoter marks and the RNA-seq data. (D-E) Peak coverage for each subset of two PC replicates, the full set of three PC replicates, and the HC duplicate (D). (E) The peak coverage ratio showed a similar coverage for the activating histone marks except H3K27ac, which was increased in PC. The repressive histone mark H3K27me3 had a higher peak coverage in HC. H3K9me3 peaks varied between replicates, activating marks (green), repressive marks (red). (F) Calculating the H3K27me3 peak coverage for each chromosome revealed increased decoration of the X chromosome, indicating an overrepresentation of female mice in the dataset. Representative chromosomes are shown. (G-H) Analysis of H3K27me3 coverage including (G) or excluding (H) peaks on gonosomes supports a gonosome-independent enrichment of H3K27me3 in HC compared to PC.

Supplementary figure 2: 15 state ChromHMM model derived from chondrocytes.

(A) Log-likelihood and (B) Akaike information criterion were calculated for ChromHMM models with increasing numbers of histone states. The minimum number of meaningful ChromHMM states was set to 15. (C) The transition probability matrix of the ChromHMM model using the epigenetic data from chondrocytes depicting the probability of state transitions. (D) Region-specific genomic enrichment of ChromHMM states for each cell type with respect to different functional annotations: Transcription start site (TSS), transcription end site (TES), zinc finger genes (ZNF-genes).

Supplementary figure 3: ChromHMM models derive from varying numbers of PC datasets and additional datasets derived from the Encode database.

(A) Comparison of 15-state ChromHMM models derived from the HC duplicate in combination with each subset of PC duplicates (M1-M3) and the PC triplicate (M0).

(B) Hierarchical clustering of genome-wide ChromHMM state vectors for the different models (M0-M3). (A-B) M1 assigned ChromHMM states identical to the states called in the original model with similar emission probabilities. The remaining two models gained one new state each. M2 gained an active state, combining the promoter marks H3K4me3, H3K9ac and the transcription-associated mark H3K36me3. M3 gained a state combing H3K36me3 with H3K9me3 and H3K27me3. Importantly, the genome-wide frequencies of these additional states and the missed state 15 from the original model were among the lowest of the different ChromHMM states. As low-frequency states can be missed when training a model with limited numbers of states, we conclude that the original model is robust. (C) The ratio of each state shows the variation between the PC subset-based models M1-M3 and the model based on the full PC triplicate M0. (D) The emission probability diagram of a 15 state ChromHMM model trained on the generalized model, representing the combined data of PC, HC and E11.5-E15.5 limb bud and brain. Chromatin states were similar to those of the model trained exclusively on chondrocyte data (compare Fig. 2A). The two states, state 15 representing a combination of H3K36me3 and H3K27me3 (repressed-active gene) and state 11, representing a combination of H3K36me3 and activating promoter marks (TSS expressed gene) were not detected, pointing to an overrepresentation of these states in chondrocytes compared to other tissues of the same developmental stage. The generalized model included a greater variety of promoter marks. Activating states (green) repressed states (red), repressed-active states (purple), empty state (gray). (E-F) Posterior predictive check of the modeled enrichment of transitions in housekeeping (E) and the chondrocyte-specific genes (F). Dots represent the measured values. 11% to 89% density of the posterior distribution (grey bars), y: number of genes with a given transition (compare Fig. 2 C-D).

Supplementary figure 4: Correlation of RNA-seq and ChIP-seq data.

(A-B) Correlation of the duplicate RNA-seq data of PC (A) and HC (B). (C) Correlation of TPM values in PC (orange) and HC (blue) and the mean state prevalence of ChromHMM states at the transcriptional start site TSS +/− 2.5 kb. High TPM values were found for activating promoter states, while repressed states were present on low expressed genes. (D-E) IGV images of the genomic regions of *Dlk1* (D) and *CD44* (E). (D) *Dlk1* carried activating marks in PC and repressed-active marks in HC. (E) *CD44* was decorated with activating marks in HC, while no enrichment of marks was detected in PC.

Supplementary figure 5: Identification of cell type specific enhancers and super-enhancers.

(A) Distribution of the proximal enhancer state 10 (red), the intragenic enhancer state 5 (blue) and the intergenic enhancer state 7 (green) with respect to their distance to the nearest gene. The intergenic state 5 and the intragenic state 7 were merged to define distal enhancers. Each enhancer state was divided into enhancer states occurring in PC, in HC, or in both cell types (common). (B) Percentage of proximal (red) and distal (green) enhancer states found specifically in PC, HC or in both (common). (C) Enrichment analysis of the genes assigned to SEs in PC, HC or common SEs. While SEs define chondrocyte-related pathways in PC, SEs were assigned to genes regulating metabolic processes in HC. (D) Posterior predictive check of the colocalization of Sox9 peaks and SEs. Empirical density distribution (dark blue line), 100 draws from the posterior (light blue lines). (E) Posterior predictive check of the calculated correlation of PC-specific, HC-specific and common SEs on differential gene expression compared to genes without a SE. Empirical density distribution (dark blue line), 100 draws from the posterior (light blue lines). (F) Enrichment of enhancer peaks in TADs. The number of H3K27ac peaks outside the TSS was compared in PC and HC. Two different TAD datasets of embryonic stem cells were used (TAD HindIII, compare figure 6C, and TAD NcoI, (30)). (G) TADs were ranked according to their differential occurrence of H3K27ac peaks. In PC, the TAD containing *Sox5* was enriched. Enrichment of the Sox9-containing TAD was not detected in the TAD (NcoI), due to differences in TAD borders. In HC a TAD which includes the *Col10a1* gene had the highest differential H3K27ac peaks.

## Supplementary tables

Supplementary table 1: Read, peak and gene number of the ChIP-seq data of the analyzed histone marks for each replicate and the conserved dataset. The calculated quality metrics for each ChIP-seq dataset are given.

Supplementary table 2: Chondrogenesis-related genes and housekeeping genes used for the state transition analysis.

Supplementary table 3: Differential expression values of chondrogenesis-related genes with the transition from (A) the activating promoter states 9 or 11 to the repressed-active state 13. blue: transition from state 9 to state 13; orange: transitions from state 11 to 13; black: genes with both transitions. The TPM and beta-values measured in RNA-seq, the published expression data, cell type and references are given for each gene. (B) Genes with the transition from the repressed state 1 to the repressed-active state 13 and vice-versa. The TPM and beta-values measured in RNA-seq are given for each gene.

Supplementary table 4: Z-scores of differential RNA-seq and ChIP-seq data used for Pareto optimization. According to their z-scores, genes were ranked into 55 Pareto fronts.

Supplementary table 5: Genes assigned to SEs in each cell type with their respective differential expression calculated as beta-values. Genes carrying SEs at different localizations in the two different cell types are denoted as shifted enhancer.

Supplementary table 6: The number of H3K27ac peaks outside a TSS (+/− 2.5 kb) within a TAD was calculated in PC and HC, respectively. TADs were ranked according to the ratio of H3K27ac peaks. Two TAD datasets of embryonic stem cells (30) were used.

## Notes

### Competing Interest Statement

The authors have declared no competing interest.

## References

1. Mikkelsen TS, Ku M, Jaffe DB, Issac B, Lieberman E, Giannoukos G, et al. Genome-wide maps of chromatin state in pluripotent and lineage-committed cells. Nature. 2007;448(7153):553–60.

2. Wu H, Gordon JA, Whitfield TW, Tai PW, van Wijnen AJ, Stein JL, et al. Chromatin dynamics regulate mesenchymal stem cell lineage specification and differentiation to osteogenesis. Biochim Biophys Acta Gene Regul Mech. 2017;1860(4):438–49.

3. Bradley EW, McGee-Lawrence ME, Westendorf JJ. Hdac-mediated control of endochondral and intramembranous ossification. Crit Rev Eukaryot Gene Expr. 2011;21(2):101–13.

4. Carpio LR, Westendorf JJ. Histone Deacetylases in Cartilage Homeostasis and Osteoarthritis. Curr Rheumatol Rep. 2016;18(8):52.

5. Kozhemyakina E, Lassar AB, Zelzer E. A pathway to bone: signaling molecules and transcription factors involved in chondrocyte development and maturation. Development. 2015;142(5):817–31.

6. Wuelling M, Vortkamp A. Chondrocyte proliferation and differentiation. Endocr Dev. 2011;21:1–11.

7. Guenther MG, Levine SS, Boyer LA, Jaenisch R, Young RA. A chromatin landmark and transcription initiation at most promoters in human cells. Cell. 2007;130(1):77–88.

8. Wagner EJ, Carpenter PB. Understanding the language of Lys36 methylation at histone H3. Nat Rev Mol Cell Biol. 2012;13(2):115–26.

9. Volpe TA, Kidner C, Hall IM, Teng G, Grewal SI, Martienssen RA. Regulation of heterochromatic silencing and histone H3 lysine-9 methylation by RNAi. Science. 2002;297(5588):1833–7.

10. Zhang T, Cooper S, Brockdorff N. The interplay of histone modifications - writers that read. EMBO Rep. 2015;16(11):1467–81.

11. Li F, Wan M, Zhang B, Peng Y, Zhou Y, Pi C, et al. Bivalent Histone Modifications and Development. Curr Stem Cell Res Ther. 2018;13(2):83–90.

12. Alder O, Lavial F, Helness A, Brookes E, Pinho S, Chandrashekran A, et al. Ring1B and Suv39h1 delineate distinct chromatin states at bivalent genes during early mouse lineage commitment. Development. 2010;137(15):2483–92.

13. Bernstein BE, Mikkelsen TS, Xie X, Kamal M, Huebert DJ, Cuff J, et al. A bivalent chromatin structure marks key developmental genes in embryonic stem cells. Cell. 2006;125(2):315–26.

14. Mauser R, Kungulovski G, Keup C, Reinhardt R, Jeltsch A. Application of dual reading domains as novel reagents in chromatin biology reveals a new H3K9me3 and H3K36me2/3 bivalent chromatin state. Epigenetics Chromatin. 2017;10(1):45.

15. Hahn MA, Wu X, Li AX, Hahn T, Pfeifer GP. Relationship between gene body DNA methylation and intragenic H3K9me3 and H3K36me3 chromatin marks. PLoS One. 2011;6(4):e18844.

16. Ohba S, He X, Hojo H, McMahon AP. Distinct Transcriptional Programs Underlie Sox9 Regulation of the Mammalian Chondrocyte. Cell Rep. 2015;12(2):229–43.

17. Liu CF, Lefebvre V. The transcription factors SOX9 and SOX5/SOX6 cooperate genome-wide through super-enhancers to drive chondrogenesis. Nucleic Acids Res. 2015;43(17):8183–203.

18. Guo M, Liu Z, Willen J, Shaw CP, Richard D, Jagoda E, et al. Epigenetic profiling of growth plate chondrocytes sheds insight into regulatory genetic variation influencing height. Elife. 2017;6.

19. Starmer J, Magnuson T. Detecting broad domains and narrow peaks in ChIP-seq data with hiddenDomains. BMC bioinformatics. 2016;17:144.

20. Lefebvre V, Li P, de Crombrugghe B. A new long form of Sox5 (L-Sox5), Sox6 and Sox9 are coexpressed in chondrogenesis and cooperatively activate the type II collagen gene. EMBO J. 1998;17(19):5718–33.

21. de Jonge HJ, Fehrmann RS, de Bont ES, Hofstra RM, Gerbens F, Kamps WA, et al. Evidence based selection of housekeeping genes. PLoS One. 2007;2(9):e898.

22. Iwamoto M, Yagami K, Lu Valle P, Olsen BR, Petropoulos CJ, Ewert DL, et al. Expression and role of c-myc in chondrocytes undergoing endochondral ossification. J Biol Chem. 1993;268(13):9645–52.

23. Yang Y, Topol L, Lee H, Wu J. Wnt5a and Wnt5b exhibit distinct activities in coordinating chondrocyte proliferation and differentiation. Development. 2003;130(5):1003–15.

24. Creyghton MP, Cheng AW, Welstead GG, Kooistra T, Carey BW, Steine EJ, et al. Histone H3K27ac separates active from poised enhancers and predicts developmental state. Proc Natl Acad Sci U S A. 2010;107(50):21931–6.

25. Whyte WA, Orlando DA, Hnisz D, Abraham BJ, Lin CY, Kagey MH, et al. Master transcription factors and mediator establish super-enhancers at key cell identity genes. Cell. 2013;153(2):307–19.

26. Tan Z, Niu B, Tsang KY, Melhado IG, Ohba S, He X, et al. Synergistic co-regulation and competition by a SOX9-GLI-FOXA phasic transcriptional network coordinate chondrocyte differentiation transitions. PLoS Genet. 2018;14(4):e1007346.

27. Lewandowski JP, Du F, Zhang S, Powell MB, Falkenstein KN, Ji H, et al. Spatiotemporal regulation of GLI target genes in the mammalian limb bud. Dev Biol. 2015;406(1):92–103.

28. Rowley MJ, Corces VG. Organizational principles of 3D genome architecture. Nat Rev Genet. 2018;19(12):789–800.

29. Hnisz D, Abraham BJ, Lee TI, Lau A, Saint-Andre V, Sigova AA, et al. Super-enhancers in the control of cell identity and disease. Cell. 2013;155(4):934–47.

30. Dixon JR, Selvaraj S, Yue F, Kim A, Li Y, Shen Y, et al. Topological domains in mammalian genomes identified by analysis of chromatin interactions. Nature. 2012;485(7398):376–80.

31. Sodersten E, Toskas K, Rraklli V, Tiklova K, Bjorklund AK, Ringner M, et al. A comprehensive map coupling histone modifications with gene regulation in adult dopaminergic and serotonergic neurons. Nat Commun. 2018;9(1):1226.

32. Yang L, Tsang KY, Tang HC, Chan D, Cheah KS. Hypertrophic chondrocytes can become osteoblasts and osteocytes in endochondral bone formation. Proc Natl Acad Sci U S A. 2014;111(33):12097–102.

33. Zhou X, von der Mark K, Henry S, Norton W, Adams H, de Crombrugghe B. Chondrocytes transdifferentiate into osteoblasts in endochondral bone during development, postnatal growth and fracture healing in mice. PLoS Genet. 2014;10(12):e1004820.

34. Tsang KY, Chan D, Cheslett D, Chan WC, So CL, Melhado IG, et al. Surviving endoplasmic reticulum stress is coupled to altered chondrocyte differentiation and function. PLoS Biol. 2007;5(3):e44.

35. Wang C, Tan Z, Niu B, Tsang KY, Tai A, Chan WCW, et al. Inhibiting the integrated stress response pathway prevents aberrant chondrocyte differentiation thereby alleviating chondrodysplasia. Elife. 2018;7.

36. Voigt P, Tee WW, Reinberg D. A double take on bivalent promoters. Genes Dev. 2013;27(12):1318–38.

37. Zhu J, Adli M, Zou JY, Verstappen G, Coyne M, Zhang X, et al. Genome-wide chromatin state transitions associated with developmental and environmental cues. Cell. 2013;152(3):642–54.

38. Kim JK, Haselgrove JC, Shapiro IM. Measurement of metabolic events in the avian epiphyseal growth cartilage using a bioluminescence technique. J Histochem Cytochem. 1993;41(5):693–702.

39. Kudelko M, Chan CW, Sharma R, Yao Q, Lau E, Chu IK, et al. Label-Free Quantitative Proteomics Reveals Survival Mechanisms Developed by Hypertrophic Chondrocytes under ER Stress. J Proteome Res. 2016;15(1):86–99.

40. Lee SY, Abel ED, Long F. Glucose metabolism induced by Bmp signaling is essential for murine skeletal development. Nat Commun. 2018;9(1):4831.

41. Chen J, Long F. mTORC1 signaling controls mammalian skeletal growth through stimulation of protein synthesis. Development. 2014;141(14):2848–54.

42. Cooper KL, Oh S, Sung Y, Dasari RR, Kirschner MW, Tabin CJ. Multiple phases of chondrocyte enlargement underlie differences in skeletal proportions. Nature. 2013;495(7441):375–8.

43. Wang J, Zhou J, Cheng CM, Kopchick JJ, Bondy CA. Evidence supporting dual, IGF-I-independent and IGF-I-dependent, roles for GH in promoting longitudinal bone growth. J Endocrinol. 2004;180(2):247–55.

44. Visel A, Akiyama JA, Shoukry M, Afzal V, Rubin EM, Pennacchio LA. Functional autonomy of distant-acting human enhancers. Genomics. 2009;93(6):509–13.

45. Pennacchio LA, Ahituv N, Moses AM, Prabhakar S, Nobrega MA, Shoukry M, et al. In vivo enhancer analysis of human conserved non-coding sequences. Nature. 2006;444(7118):499–502.

46. Yu M, Ren B. The Three-Dimensional Organization of Mammalian Genomes. Annu Rev Cell Dev Biol. 2017;33:265–89.

47. Moorthy SD, Davidson S, Shchuka VM, Singh G, Malek-Gilani N, Langroudi L, et al. Enhancers and super-enhancers have an equivalent regulatory role in embryonic stem cells through regulation of single or multiple genes. Genome Res. 2017;27(2):246–58.

48. Osterwalder M, Barozzi I, Tissieres V, Fukuda-Yuzawa Y, Mannion BJ, Afzal SY, et al. Enhancer redundancy provides phenotypic robustness in mammalian development. Nature. 2018;554(7691):239–43.

49. Furlong EEM, Levine M. Developmental enhancers and chromosome topology. Science. 2018;361(6409):1341–5.

50. Heinz S, Romanoski CE, Benner C, Glass CK. The selection and function of cell type-specific enhancers. Nat Rev Mol Cell Biol. 2015;16(3):144–54.

51. Terpstra L, Prud’homme J, Arabian A, Takeda S, Karsenty G, Dedhar S, et al. Reduced chondrocyte proliferation and chondrodysplasia in mice lacking the integrin-linked kinase in chondrocytes. J Cell Biol. 2003;162(1):139–48.

52. Gebhard S, Hattori T, Bauer E, Schlund B, Bosl MR, de Crombrugghe B, et al. Specific expression of Cre recombinase in hypertrophic cartilage under the control of a BAC-Col10a1 promoter. Matrix Biol. 2008;27(8):693–9.

53. Srinivas S, Watanabe T, Lin CS, William CM, Tanabe Y, Jessell TM, et al. Cre reporter strains produced by targeted insertion of EYFP and ECFP into the ROSA26 locus. BMC Dev Biol. 2001;1:4.

54. Kharchenko PV, Tolstorukov MY, Park PJ. Design and analysis of ChIP-seq experiments for DNA-binding proteins. Nature biotechnology. 2008;26:1351–9.

55. Landt SG, Marinov GK, Kundaje A, Kheradpour P, Pauli F, Batzoglou S, et al. ChIP-seq guidelines and practices of the ENCODE and modENCODE consortia. Genome Res. 2012;22(9):1813–31.

56. Robinson JT, Thorvaldsdottir H, Winckler W, Guttman M, Lander ES, Getz G, et al. Integrative genomics viewer. Nat Biotechnol. 2011;29(1):24–6.

57. Ernst J, Kellis M. ChromHMM: automating chromatin-state discovery and characterization. Nat Methods. 2012;9(3):215–6.

58. Lawrence M, Huber W, Pagès H, Aboyoun P, Carlson M, Gentleman R, et al. Software for computing and annotating genomic ranges. PLoS computational biology. 2013;9:e1003118.

59. Burkner PC. Advanced Bayesian Multilevel Modeling with the R Package brms. R J. 2018;10(1):395–411.

60. Huang J, Li K, Cai W, Liu X, Zhang Y, Orkin SH, et al. Dissecting super-enhancer hierarchy based on chromatin interactions. Nat Commun. 2018;9(1):943.

61. McLean CY, Bristor D, Hiller M, Clarke SL, Schaar BT, Lowe CB, et al. GREAT improves functional interpretation of cis-regulatory regions. Nat Biotechnol. 2010;28(5):495–501.

62. Hung JH, Weng Z. Mapping Short Sequence Reads to a Reference Genome. Cold Spring Harb Protoc. 2017;2017(2).

63. Bolger AM, Lohse M, Usadel B. Trimmomatic: a flexible trimmer for Illumina sequence data. Bioinformatics. 2014;30(15):2114–20.

64. Bray NL, Pimentel H, Melsted P, Pachter L. Near-optimal probabilistic RNA-seq quantification. Nat Biotechnol. 2016;34(5):525–7.

65. Pimentel H, Bray NL, Puente S, Melsted P, Pachter L. Differential analysis of RNA-seq incorporating quantification uncertainty. Nat Methods. 2017;14(7):687–90.

66. Yu G, Wang LG, Han Y, He QY. clusterProfiler: an R package for comparing biological themes among gene clusters. OMICS. 2012;16(5):284–7.

67. Shwartz Y, Zelzer E. Nonradioactive in situ hybridization on skeletal tissue sections. Methods Mol Biol. 2014;1130:203–15.

68. Bachvarova V, Dierker T, Esko J, Hoffmann D, Kjellen L, Vortkamp A. Chondrocytes respond to an altered heparan sulfate composition with distinct changes of heparan sulfate structure and increased levels of chondroitin sulfate. Matrix Biol. 2020.

69. Li H, Durbin R. Fast and accurate short read alignment with Burrows-Wheeler transform. Bioinformatics (Oxford, England). 2009;25:1754–60.

70. Li H, Handsaker B, Wysoker A, Fennell T, Ruan J, Homer N, et al. The Sequence Alignment/Map format and SAMtools. Bioinformatics (Oxford, England). 2009;25:2078–9.

71. Love MI, Huber W, Anders S. Moderated estimation of fold change and dispersion for RNA-seq data with DESeq2. Genome Biol. 2014;15(12):550.

