## Supplementay Figurese for "Epigenetic mechanisms mediating cell state transitions in chondrocytes"

### Supplementary Figure 1

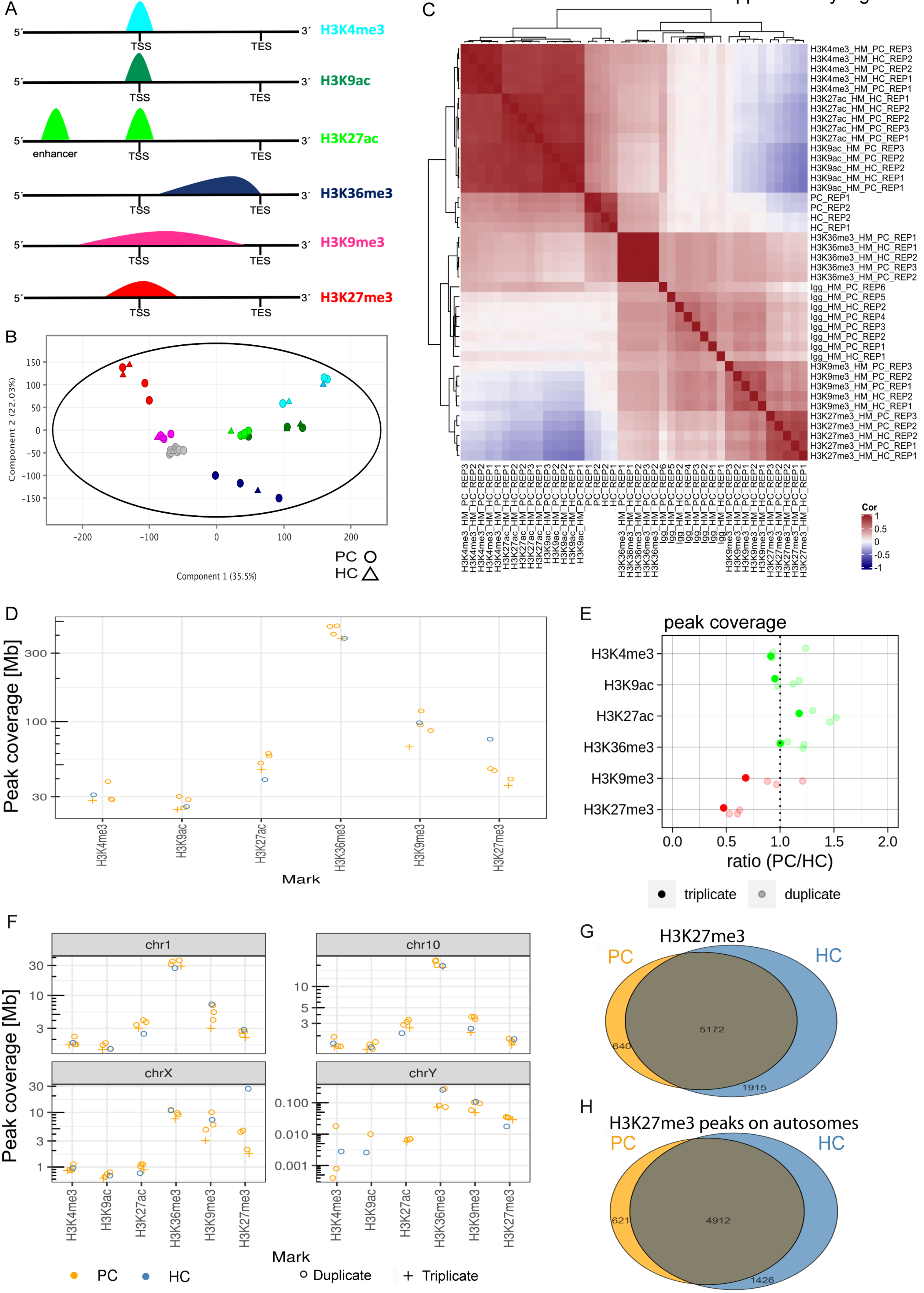

Supplementary Figure 2

A

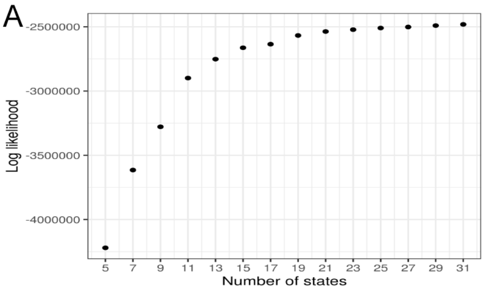


B

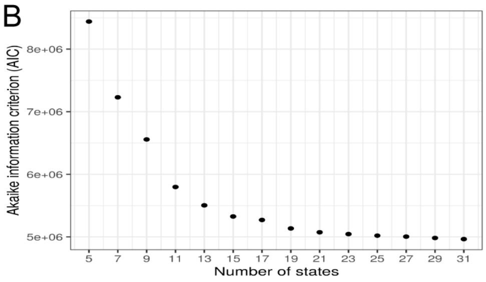


C

| State (from) \ State (to) | 1 | 2 | 3 | 4 | 5 | 6 | 7 | 8 | 9 | 10 | 11 | 12 | 13 | 14 | 15 |
|---|---|---|---|---|---|---|---|---|---|---|---|---|---|---|---|
| 15 | 0 | 0 | 0.005 | 0.027 | 0 | 0.007 | 0 | 0 | 0 | 0 | 0 | 0 | 0.008 | 0.034 | 0.918 |
| 14 | 0.005 | 0 | 0 | 0 | 0 | 0.03 | 0 | 0 | 0 | 0 | 0 | 0 | 0.015 | 0.948 | 0.002 |
| 13 | 0.013 | 0 | 0 | 0 | 0 | 0.003 | 0.001 | 0.002 | 0.007 | 0 | 0.001 | 0.005 | 0.861 | 0.103 | 0.004 |
| 12 | 0 | 0 | 0 | 0.007 | 0 | 0.262 | 0.001 | 0.027 | 0.168 | 0.003 | 0.007 | 0.517 | 0.007 | 0.001 | 0 |
| 11 | 0 | 0 | 0.002 | 0.077 | 0.032 | 0 | 0 | 0.002 | 0.039 | 0.007 | 0.833 | 0.007 | 0.001 | 0 | 0 |
| 10 | 0 | 0 | 0 | 0.006 | 0.004 | 0.133 | 0.129 | 0 | 0.204 | 0.507 | 0.013 | 0.005 | 0.001 | 0 | 0 |
| 9 | 0 | 0 | 0 | 0 | 0 | 0.015 | 0.015 | 0.027 | 0.859 | 0.025 | 0.01 | 0.045 | 0.003 | 0 | 0 |
| 8 | 0 | 0 | 0 | 0.002 | 0.002 | 0.127 | 0.145 | 0.38 | 0.261 | 0.001 | 0.004 | 0.07 | 0.006 | 0 | 0 |
| 7 | 0 | 0 | 0 | 0.003 | 0.01 | 0.141 | 0.794 | 0.016 | 0.017 | 0.017 | 0 | 0 | 0.001 | 0 | 0 |
| 6 | 0 | 0.001 | 0 | 0.003 | 0 | 0.993 | 0.001 | 0 | 0 | 0 | 0 | 0.001 | 0 | 0.001 | 0 |
| 5 | 0 | 0 | 0 | 0.128 | 0.823 | 0.002 | 0.023 | 0 | 0 | 0.001 | 0.021 | 0 | 0 | 0 | 0 |
| 4 | 0 | 0 | 0.002 | 0.975 | 0.002 | 0.019 | 0 | 0 | 0 | 0 | 0.001 | 0 | 0 | 0 | 0 |
| 3 | 0.001 | 0.008 | 0.919 | 0.055 | 0 | 0.015 | 0 | 0 | 0 | 0 | 0 | 0 | 0 | 0 | 0.001 |
| 2 | 0.001 | 0.951 | 0.002 | 0 | 0 | 0.046 | 0 | 0 | 0 | 0 | 0 | 0 | 0 | 0 | 0 |
| 1 | 0.95 | 0.007 | 0.002 | 0 | 0 | 0.005 | 0 | 0 | 0 | 0 | 0 | 0 | 0.009 | 0.026 | 0 |

P: 1.00, 0.75, 0.50, 0.25, 0.00

D

PC

| | State | RefSeqGene | RefSeqExon | RefSeqTSS | RefSeqTES | RefSeqTSS2kb | CpGIsland | Vista | Fantom all | Fantom forelimb | Zfp |
|---|---|---|---|---|---|---|---|---|---|---|---|
| promoter | 15 | 1.76 | 5.81 | 1.93 | 6.31 | 3.93 | 11.27 | 0.65 | 1.41 | 2.02 | 3.89 |
| promoter | 14 | 1.19 | 2.13 | 2.66 | 2.56 | 6.46 | 2.44 | 2.71 | 2.71 | 1.58 | 0.67 |
| primed promoter | 13 | 1.84 | 9.47 | 43.89 | 3.18 | 19.73 | 99.71 | 3.36 | 6.64 | 16.88 | 2.37 |
| promoter | 12 | 1.65 | 4.64 | 30.58 | 2.18 | 19.12 | 27.04 | 4.4 | 7.5 | 19.97 | 3.39 |
| active promoter | 11 | 2.45 | 7.92 | 19.62 | 8.24 | 16.64 | 29.85 | 8.27 | 6.23 | 23.95 | 4.7 |
| proximal enhancer | 10 | 1.61 | 2.2 | 6.18 | 3.01 | 16.04 | 4.43 | 29.66 | 11.52 | 39.05 | 2.8 |
| intergenic enhancer | 9 | 2.02 | 8.83 | 62.34 | 2.1 | 24.31 | 95.47 | 5.77 | 6.54 | 38.13 | 4.25 |
| empty | 8 | 1.51 | 4.17 | 32.43 | 2.25 | 17.76 | 29.36 | 12.04 | 12.24 | 43.73 | 2.66 |
| intragenic enhancer | 7 | 1.19 | 0.81 | 1.71 | 1.05 | 3.57 | 0.48 | 22.04 | 7.99 | 12.39 | 0.89 |
| expressed gene | 6 | 0.79 | 0.54 | 0.44 | 0.68 | 0.69 | 0.05 | 0.71 | 0.81 | 0.42 | 0.56 |
| ZNF-gene | 5 | 2.17 | 1.54 | 0.73 | 1.48 | 1.45 | 0.32 | 15.71 | 5.53 | 10.28 | 2.63 |
| heterochromatin | 4 | 2.26 | 3.16 | 0.33 | 2.59 | 0.55 | 0.21 | 0.95 | 1.17 | 1.22 | 2.81 |
| silenced gene | 3 | 1.36 | 1.97 | 0.4 | 2.15 | 0.61 | 1.18 | 0 | 0.55 | 1.04 | 16.41 |
| repression | 2 | 0.43 | 0.52 | 0.81 | 0.93 | 0.89 | 0.15 | 0.03 | 0.2 | 0.12 | 1.46 |
| repressed-active promotor | 1 | 1.28 | 4.28 | 9 | 4.5 | 8.69 | 28.45 | 5.74 | 4.08 | 3.82 | 1.48 |

HC

| State | RefSeqGene | RefSeqExon | RefSeqTSS | RefSeqTES | RefSeqTSS2kb | CpGIsland | Vista | Fantom all | Fantom forelimb | Zfp |
|---|---|---|---|---|---|---|---|---|---|---|
| 15 | 2.21 | 4.83 | 1.15 | 4.46 | 1.92 | 4.2 | 0.28 | 1.17 | 4.13 | 4.71 |
| 14 | 1.06 | 1.38 | 1.54 | 1.65 | 3.5 | 1.01 | 2.05 | 1.51 | 0.88 | 0.44 |
| 13 | 1.83 | 8.67 | 40.87 | 2.57 | 19.6 | 89.7 | 4.76 | 7 | 20.22 | 1.71 |
| 12 | 1.66 | 4.65 | 31.48 | 2.28 | 19.56 | 27.32 | 5.78 | 7.94 | 24.42 | 3.82 |
| 11 | 2.45 | 8.12 | 24.6 | 7.52 | 18.65 | 33.38 | 7.2 | 6.07 | 22 | 4.38 |
| 10 | 1.74 | 2.27 | 6.16 | 3.08 | 17.46 | 2.69 | 27.58 | 13.31 | 34.23 | 3.07 |
| 9 | 2 | 8.56 | 61.22 | 1.98 | 24.32 | 94.19 | 5.61 | 6.93 | 38.93 | 4.09 |
| 8 | 1.43 | 3.05 | 21.94 | 1.74 | 17.45 | 18.34 | 17.32 | 14.97 | 53.82 | 2.1 |
| 7 | 1.16 | 0.76 | 1.89 | 0.99 | 4.41 | 0.26 | 22.96 | 11.56 | 17.98 | 0.92 |
| 6 | 0.76 | 0.48 | 0.42 | 0.61 | 0.65 | 0.05 | 0.74 | 0.79 | 0.38 | 0.54 |
| 5 | 2.36 | 1.67 | 0.84 | 2.17 | 2.1 | 0.2 | 19.6 | 9.25 | 16.45 | 2.59 |
| 4 | 2.42 | 3.5 | 0.36 | 2.95 | 0.63 | 0.21 | 1.04 | 1.27 | 1.31 | 2.85 |
| 3 | 1.48 | 2.27 | 0.41 | 2.35 | 0.61 | 1.07 | 0.42 | 0.5 | 0.7 | 20.39 |
| 2 | 0.38 | 0.42 | 0.68 | 0.81 | 0.77 | 0.07 | 0.01 | 0.18 | 0.08 | 0.98 |
| 1 | 1.13 | 3.35 | 5.91 | 3.98 | 6.58 | 19.14 | 4.48 | 3.64 | 2.83 | 1.95 |

Fold enrichment: 0, 25, 50, 75

Supplementary Figure 3

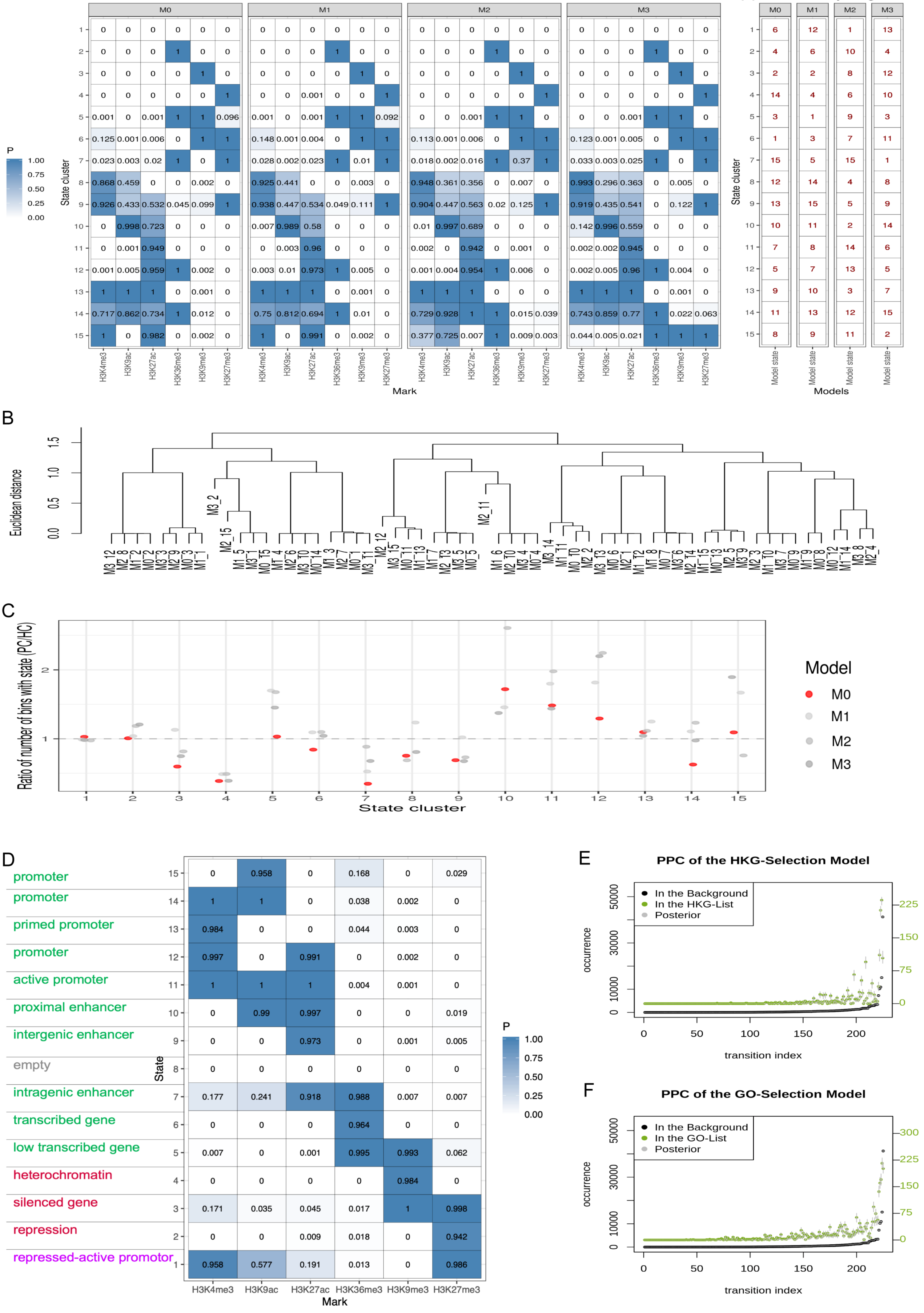

Supplementary Figure 4

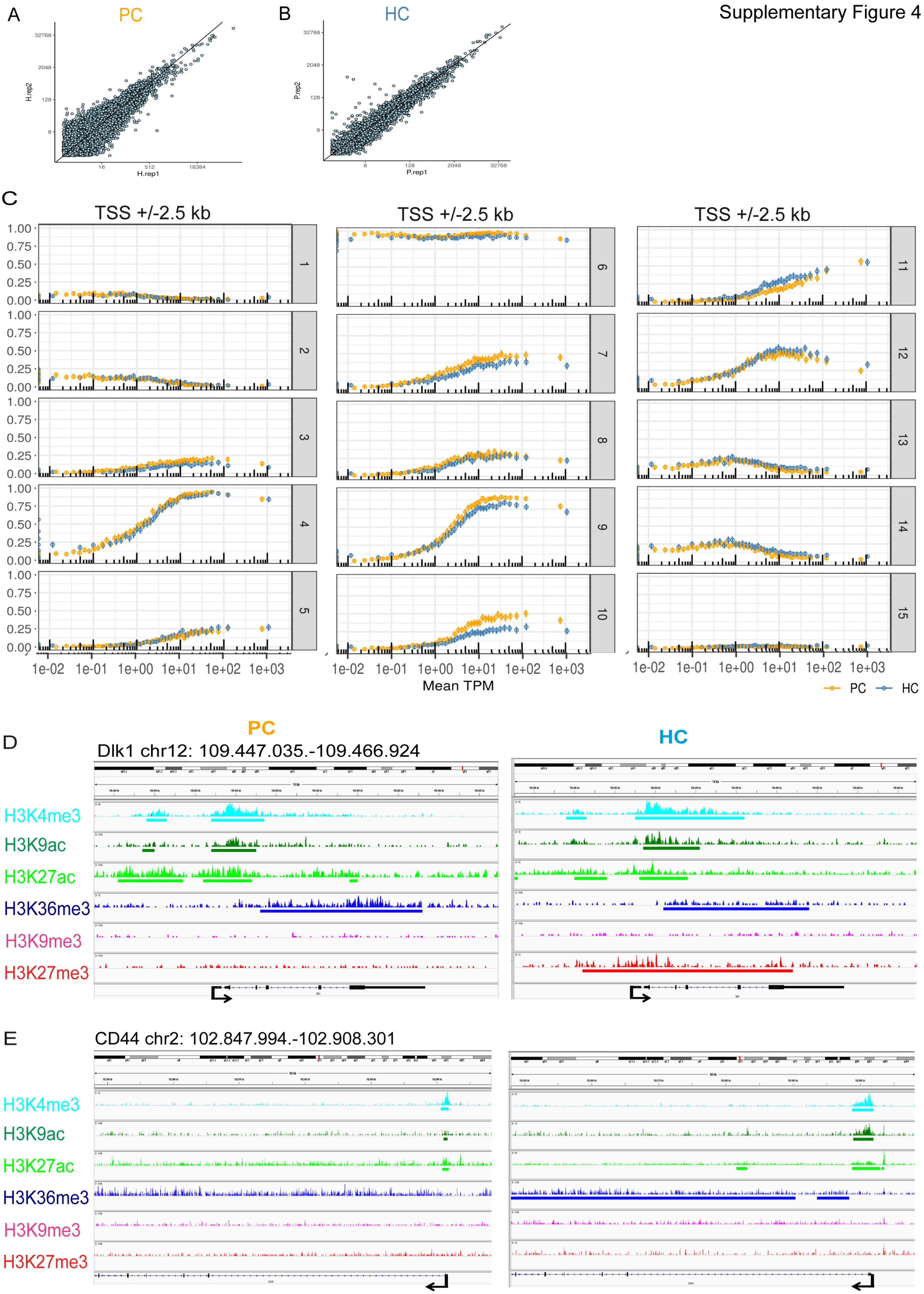

### Supplementary Figure 5

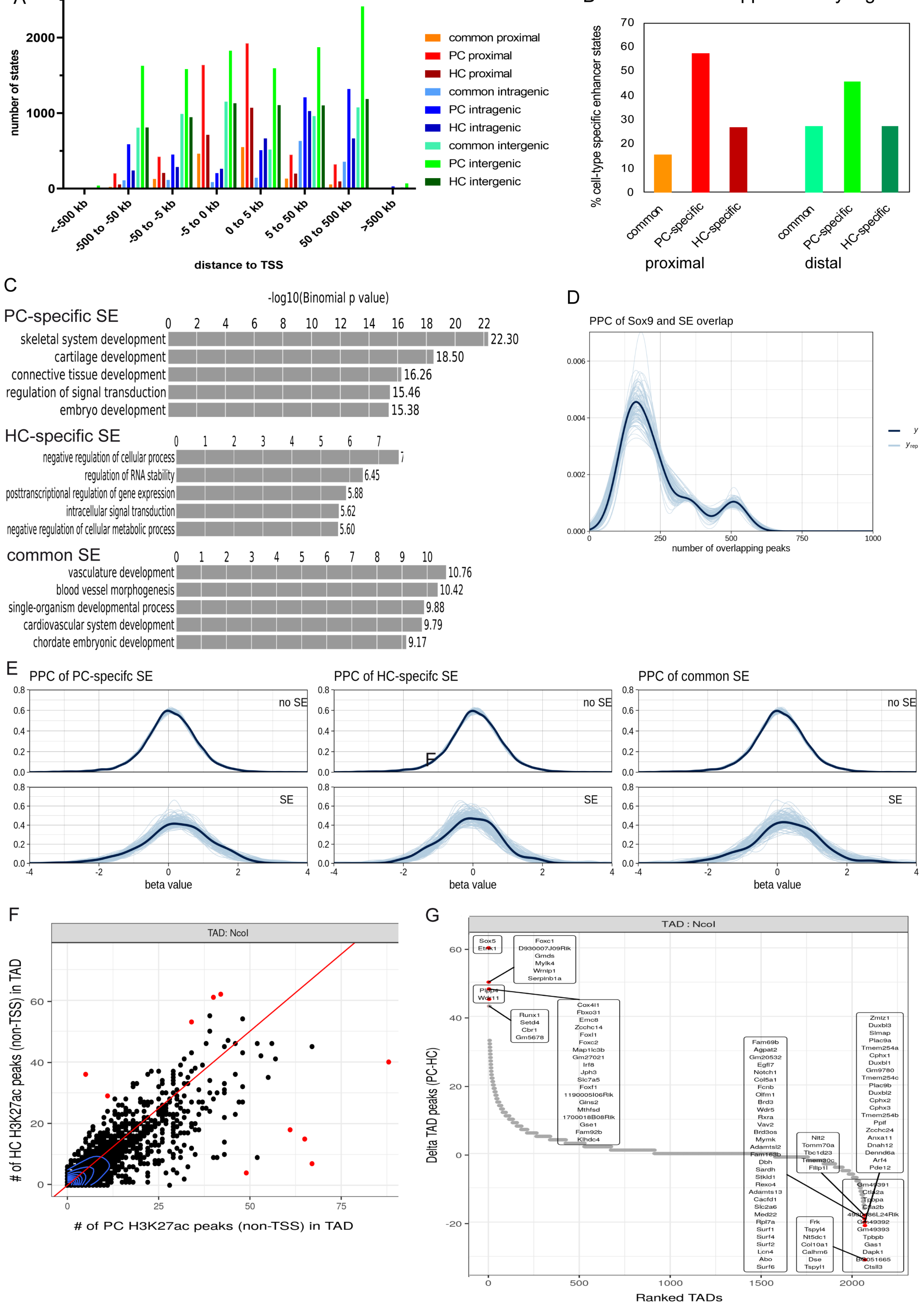
